# A Stapled Peptide Inhibitor of METTL3-METTL14 for Cancer Therapy

**DOI:** 10.1101/2023.09.04.556216

**Authors:** Zenghui Li, Yuqing Feng, Hong Han, Xingyue Jiang, Weiyu Chen, Xuezhen Ma, Yang Mei, Dan Yuan, Dingxiao Zhang, Junfeng Shi

## Abstract

METTL3, a primary methyltransferase catalyzing RNA N6-methyladenosine (m6A) modification, has been identified as an oncogene in several cancer types and thus nominated as a potentially effective target for therapeutic inhibition, although current options using this strategy are limited. In this study, we targeted protein-protein interactions at the METTL3-METTL14 binding interface to inhibit complex formation and subsequent catalysis of RNA m6A modification. Among candidate peptides, **RM3** exhibited the highest anti-cancer potency, inhibiting METTL3 activity while also facilitating its proteasomal degradation. We then designed a stapled peptide inhibitor (**RSM3**) with enhanced peptide stability and formation of the α-helical secondary structure required for METTL3 interaction. Functional and transcriptomic analysis *in vivo* indicated that **RSM3** induced upregulation of programmed cell death-related genes while inhibiting cancer-promoting signals. Furthermore, tumor growth was significantly suppressed while apoptosis was enhanced upon **RSM3** treatment, accompanied by in-creased METTL3 degradation, and reduced global RNA methylation levels in two *in vivo* tumor models. This peptide inhibitor thus exploits a mechanism distinct from other competitive-binding small molecules to inhibit oncogenic METTL3 activity. Our findings collectively highlight the potential of targeting METTL3 in cancer therapies through peptide-based inhibition of complex formation and proteolytic degradation.

## Introduction

N6-methyladenosine (m6A) is a prevalent RNA modification that plays essential regulatory roles to coordinate a wide range of biological processes, including RNA splicing, stability, and translation^[1]^. Dysregulation of m6A modification has been firmly linked to the development and progression of various diseases (*e*.*g*., cancers)^[2]^. Installation of m6A modifications requires the methyltransferase activity catalyzed by the METTL3-METTL14 complex, the former protein serving as the catalytic subunit, while the latter performs RNA-binding function; by contrast, m6A removal is mediated by m6A demethylases, such as FTO and ALKBH5^[3]^. Complex formation is essential for m6A methyltransferase function, and m6A modification is largely lost in the absence of METTL3-METTL14 heterodimerization^[4]^. Numerous studies have shown that METTL3 is expressed at aberrantly high levels in several cancer types, suggesting a prominent role in cancer progression^[5]^, and has been shown to positively regulate cancer development by promoting metastasis and tumor growth in malignancies such as acute leukemia^[6]^, liver cancer^[7]^, stomach cancer^[8]^, lung cancer^[9]^, colorectal cancer^[10]^, and prostate cancer (PCa)^[11]^. Conversely, other studies have shown that cancer cell growth is dramatically inhibited upon METTL3 knockdown, suggesting its potential as a therapeutic target for treating various cancers ^[12]^.

Although the crystal structure of the METTL3-METTL14 complex has been solved since 2016^[13]^, progress in developing METTL3 inhibitors has been limited, with the METTL3 substrate co-factor, S-Adenyl-methionine (SAM), which is required for methyl group transfer, serving as the most common target for inhibitor design. For example, Kouzarides and co-workers identified the small molecule METTL3 inhibitor, **STM2457**, which displayed high potency against acute myeloid leukemia in high-throughput screens, but still lacks experimental validation of its efficacy against solid tumors^[14]^. Although other competitive and non-competitive inhibitors that are structurally similar to SAM, such as Sinefungin^[15]^, UZH2^[16]^, and Eltrombopag^[17]^, have been shown to exhibit substantial suppression of cancer cell growth, the *in vivo* efficacy of these small molecules requires further investigation. Thus, METTL3 is well-established as a viable potential target for cancer therapies, but the lack of available options in clinic suggests an urgent need for further development of effective METTL3-targeting treatment strategies ^[18]^.

As an alternative to small molecules, peptides offer a versatile and potentially powerful approach for targeted inhibition of proteins, especially through competitive disruption of protein-protein interactions (PPIs) by mimicking peptide binding epitopes^[19]^. Peptide drugs have been extensively developed for several diseases such as diabetes, and show high potential for clinical application^[20]^. Here, in this work, we developed a series of peptide-based inhibitors targeting METTL3 through a rational design strategy with subsequent modification to enhance specific properties. Among the candidates, one peptide targeting a binding site on METTL3 required for complex assembly with METTL14 showed the highest potency against a panel of cancer cells, leading to our subsequent engineering of its affinity and stability, especially through Bis(bromomethyl)biphenyl (Bph)-based stapling. This final peptide inhibitor, **RMS3**, exhibited high efficacy in inhibiting the growth of multiple cancer cell lines, including PCa (*e*.*g*. PC3 and DU145), leukemia (*e*.*g*. CCRF-CEM), and lung cancer (*e*.*g*. A549). Furthermore, **RMS3** could both disrupt METTL3-METTL14 complex formation as well as induce its subsequent proteasomal degradation, globally inhibiting m6A modification in cancer cells. As the first-in-class peptide inhibitor targeting METTL3, **RMS3** utilizes a markedly distinct mechanism of action from that of small molecule inhibitors, supported by *in vitro* and *in vivo* validation of its cytotoxicity and anti-tumor effects in multiple cancer cell lines and xenograft tumor models. Our study thus provides a promising therapeutic approach to treat a variety of tumor types, particularly PCa, as well as a design strategy for developing new peptide-based drugs for other METTL3-dependent diseases or other therapeutic targets.

## Results and Discussion

### Peptide inhibitor design and validation

Using previously reported crystal structure of the METTL3-METTL14 complex (PDB: 5IL1)^[13]^, we identified two distinct domains potentially essential for protein-protein interactions (PPIs) required for complex formation, suggesting their strong potential as targets for peptide-based inhibition of METTL3 activity. In the crystallized structure, METTL3 and METTL14 form tight binding interfaces through two regions: interface 1, consisting of the interface loop (residues 462-479) of METTL3, and interface 2, involving helix α2 (residues 435-447) of METTL3. Additionally, a potential target site resides in the catalytic core, with loop residues 534-541 (R536, H538, and N539) playing a pivotal role in interactions with S-adenosylmethionine (SAM)^[13]^. Building on this structural information, the **RM1, RM2**, and **RM3** candidate peptides were respectively designed to origin from the interface loop (**M1**, residues 462-479), **M2** (catalytic core, residues 534-541), and helix α2 (**M3**, residues 435-447) of the interface 2 in the complex, and fused each peptide to the classic cell penetrating peptide (CPP), R9 (RRRRRRRRR), to enhance cellular penetration (**Fig. 1 & Fig.S1-3**)^[21]^. Among the functional regions targeted for interference, **RM1** is predicted to disrupt interactions between METTL3-METTLE14 and mRNA, incorporating sequence from the RNA-binding grooves; **RM2** was designed to interact with the methylase catalytic core which binds to the SAM catalytic substrate; and **RM3** was derived from an α-helix located at the METTL3-METTL14 interface and intended to destabilize complex formation and block protein function. The detailed molecular mechanism(s) and regulatory network (METTL3/m6A/RRBP1) of METTL3 in promoting cancer progress in prostate cancer (PCa) is described in a parallel study^[22]^. Considering that METTL3 is highly upregulated in PCa cells and plays a significant role in cancer development, we assessed the potency of peptide inhibitors using MTT assays with PC3 and DU145 cell lines. As shown in **Fig. 2a and 2b, RM1** and **RM2** exhibited relatively modest cytotoxicity, while **RM3** displayed substantially greater anticancer effects, with IC_50_ values of 35.4 μM and 55.3 μM against PC3 and DU145 cells, respectively. Further evaluation through long-term clonogenic assays (**Fig. 2c and 2d**) confirmed that **RM3** exerted stronger inhibitory effects than either **RM1** or **RM2**. These findings supported the further exploration of **RM3** as a promising candidate anticancer drug warranting further optimization.

**Figure 1.**
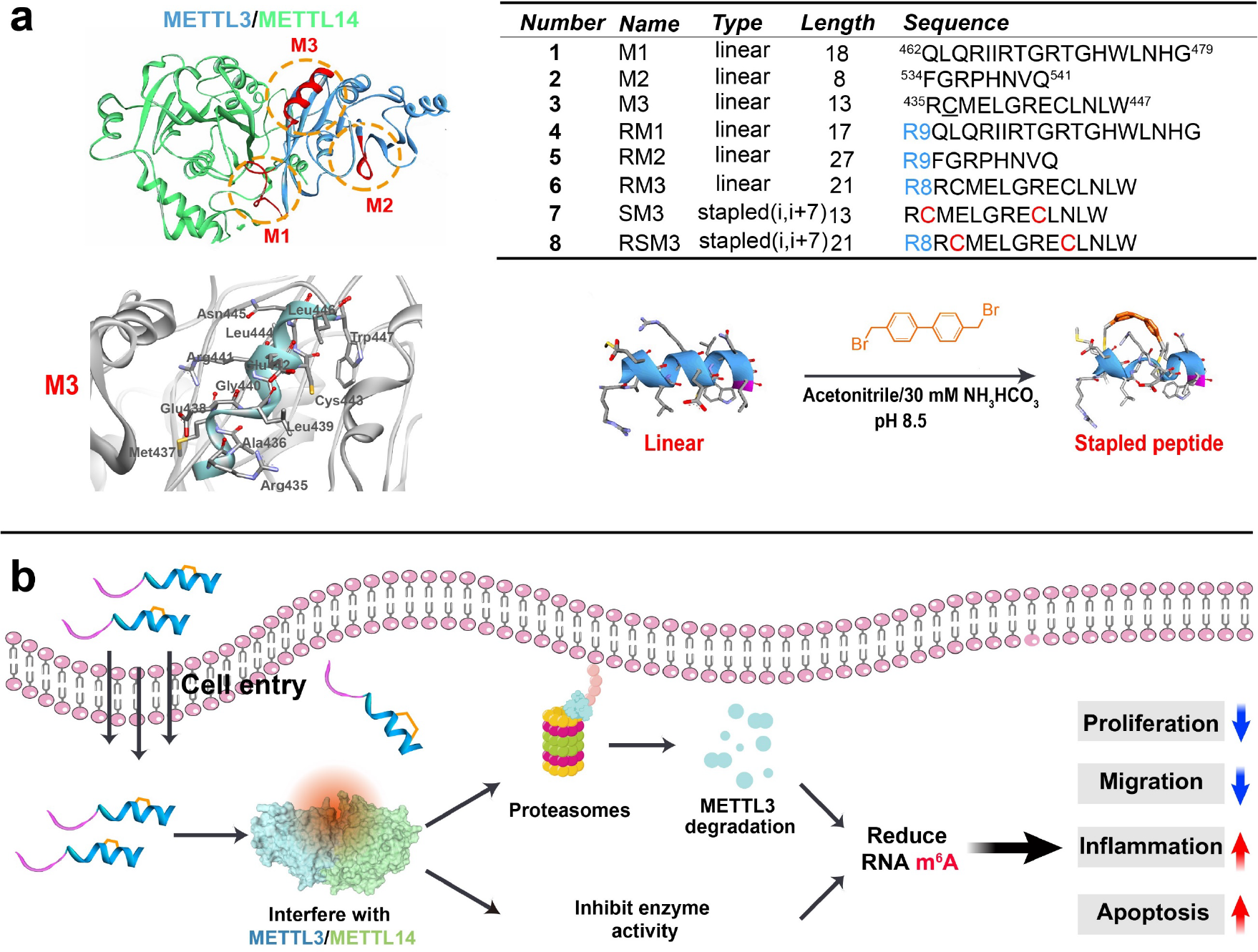
**(a)** Model of the METTL3-METTL14 complex crystal structure illustrating PPIs (yellow dashed circles) at the binding interface and the peptide sequences designed to target and disrupt these PPIs in **M1, M2**, and **M3** regions. **(b)** Further optimization through peptide stapling technology enabled construction of **RMS3**, increasing the potency of peptide inhibitory effects on cancer cell proliferation, migration, and tumor growth. Note: The amino acids highlighted (M3, RCMELGRECLNLW) with underlining, which underwent mutation from Alanine to cysteine in the original sequence, were used to construct stapled peptides.

**Figure 2.**
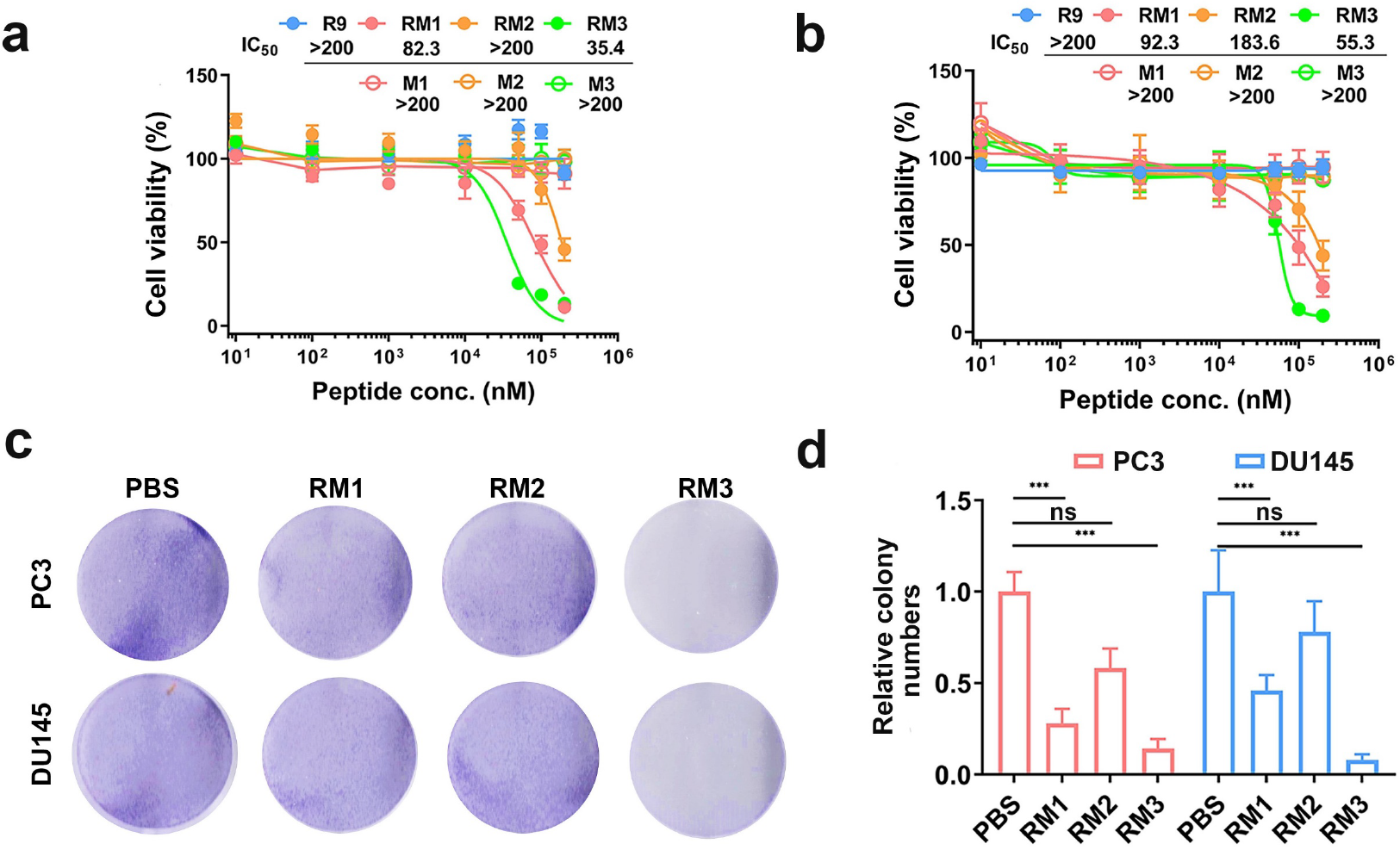
Evaluation of peptide inhibitors. Cytotoxicity of peptide inhibitors against the **(a)** PC3 and **(b)** Du145 cell lines. Clone formation plate assays **(c)** and quantification **(d)** to assess the peptide effects on growth and proliferation in PCa cells treated with 15 μM peptide inhibitor. For a-d mean and error (±s.d.) were obtained from at least three replicates. One-way analysis of variance (ANOVA) with Tukey’s test correction was used in statistical analysis test. ^*^P < 0.05, ^**^P < 0.01, ^***^P < 0.005, and not significant by P > 0.05.

### Physicochemical properties and activity of M3 derivatives in vitro

To enhance the effects of the **M3** peptide, we employed peptide stapling, a well-established technique for increasing peptide stability and activity by covalently linking two specific amino acids (e.g. Cysteine). In this work, we utilized this strategy to generate the **SM3** and **RSM3** stapled peptides from **M3** and **RM3**, respectively. Through the dynamic simulations and MMGBSA binding energy analysis of residues, it was observed that Ala436 and Cys443 are rarely involved in interactions with target proteins, as depicted in **Fig. S4a**. Utilizing visualization software (PyMOL), the distance between Ala436 and Cys443 was determined to be 12.2 Å, closely aligning with the distance of the (i+7) stapled linker Bph, as shown in **Fig. S4b**. To facilitate Cys-Cys stapling, Ala 436 was mutated to Cys due to its non-bulky and chemical inert nature^[23]^, and a crosslink was initiated using Bph with Cys443 to generate a (i+7) stapled peptide, (refer to synthetic routes in **Fig. S5**). The selection of the stapled site and design aimed to minimize potential steric clashes that could interfere with binding events while preserving peptide conformation. The binding affinity of stapled peptides was evaluated through microscale thermophoresis (MST) assays, which measure interaction strength between peptides and their target protein (**Fig. 3a**). Notably, **RSM3** exhibited high affinity binding to the METTL3-METTL14 complex, with a Kd value of 3.10 μM. In contrast, the non-stapled peptide, **RM3**, displayed significantly lower affinity (Kd = 68.25 μM). The affinity of peptide was further confirmed using Octet® BLI system (**Fig. S6**). This significant improvement in binding affinity was likely due to the higher stability secondary structure of the stapled peptide. To assess the cellular interaction of the peptide with METTL3, we conducted a biotinylated peptide pull-down assay. The results clearly indicated that the successful pulled down of the proteins METTL3 and METTL14 in PC3 cells by our peptide. This outcome serves as compelling evidence for the peptide’s ability to effectively bind to the target at the cellular level (**Fig. S7**). Circular dichroism (CD) measurements further verified that **RSM3** could adopt an α-helix conformation in solution, whereas the non-stapled **RM3** peptide fragment was mainly found in a random coil conformation (**Fig. 3b**). Functional assays demonstrated that **RSM3** could effectively inhibit METTL3-METTL14 complex transferase activity. More specifically, METTL3 methyltransferase activity was decreased by 45.5% in the presence of **RSM3**, but decreased by 34.5% upon exposure to **RM3**, further indicating the higher potency of the stapled peptide (**Fig. 3c**). Stability assays indicated that **RSM3** exhibited substantially higher biostability when treated with a 25% fetal bovine serum (FBS) solution, retaining ∼41.8% of its initial concentration after 360 min of incubation, while the **RM3** linear peptide was completely degraded within 180 min (**Fig. 3d**). This stability could be attributed to the structural integrity conferred by the stapling strategy. In addition, we evaluated possible hemolytic activity by the peptide to assess its potential cytotoxicity towards human blood cells. In primary human blood cells, the peptide inhibitor showed no obvious toxicity at concentrations up to 200 μM after 24 h of incubation (**Fig. 3e)**. Collectively, these findings highlight the improved binding affinity, biostability, and activity that are inherently linked to a highly stable α-helical conformation resulting from peptide stapling. We therefore further explored the development of **RSM3** as a promising anti-cancer drug candidate.

**Figure 3.**
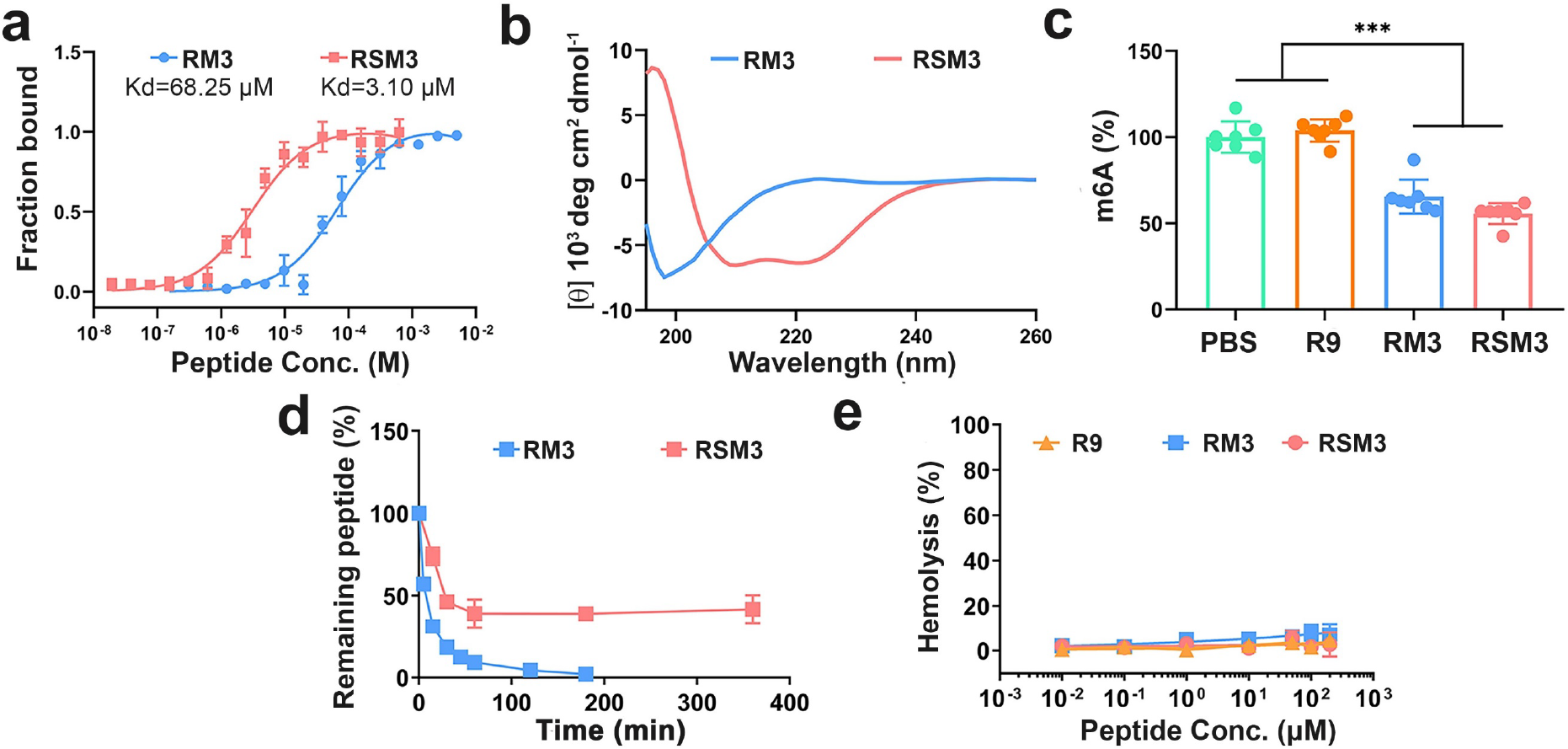
Physicochemical characterization of peptide inhibitors. **(a)** MST assays measuring the binding affinity between peptide inhibitors and METTL3–METTL14 complex, and **(b)** circular dichroism (CD) spectra scanning of **RSM3** (red) and **RM3** (blue) to determine peptide secondary structure. **(c)** Peptide effects on METTL3–METTL14 methyltransferase activity *in vitro*. **(d)** Serum stability of **RSM3** and **RM3** over time in the presence of 25% FBS. **(E)** Hemolysis assays with primary human blood cells evaluating the hemolytic potential of **R9, RM3** and **RSM3**. For **a, c-e** mean and error (±s.d.) were obtained from at least three replicates. One-way analysis of variance (ANOVA) with Tukey’s test correction was used in statistical analysis test. ^*^P < 0.05, ^**^P < 0.01, ^***^P < 0.005, and not significant by P > 0.05.

### Evaluation of RSM3 anti-cancer effects

To evaluate the therapeutic or anti-cancer effects of the **RSM3** stapled peptide, its general cytotoxicity was assessed in a panel of ten cancer cell lines. According to previous reports, METTL3 function as tumor-promoting factors in many cancer types or METTL3 inhibiting hinders tumor growth cancer growth, etc. liver cancer (HepG2) and cervical cancer (HeLa)^[24]^; colorectal cancer (HCT116)^[25]^; leucocythemia (CCRF-CEM, K562, MDSL, HL60)^[14]^, PCa cell(DU145, PC3)^[26]^.The results showed that both **RSM3** and **RM3** exhibited broad toxicity towards various cancer cells including PC3, DU145, HeLa, HepG2, HCT116, A549, K562, MDS-L, CCRF-CEM (**Fig. S8**). It is noteworthy that the IC_50_ of **RSM3** towards the leukemia cell line, CCRF-CEM, was approximately 4.6 ± 1.03 μM. Following our initial experiments, further investigation of **RSM3** activity focused on the PCa models. As expected, the **RSM3** stapled peptide displayed higher toxicity, with lower IC_50_ values, than the non-stapled **RM3** peptide (25.8 ± 2.4 μM *vs* 55.3 ± 1.9 μM, **RSM3** *vs* **RM3**) in the DU145 cell line (**Fig. 4a& Table S1**). Cell migration and colony formation assays showed the potent inhibitory effects of **RSM3** and **RM3** on cancer cell aggressiveness compared to the control peptide R9. Treatment with either 15 μM **RSM3** or **RM3** resulted in approximately 2∼3 fold less cell migration and a ∼20-fold reduction in PC3 or DU145 colony formation **(Fig. 4b&c)**.

**Figure 4.**
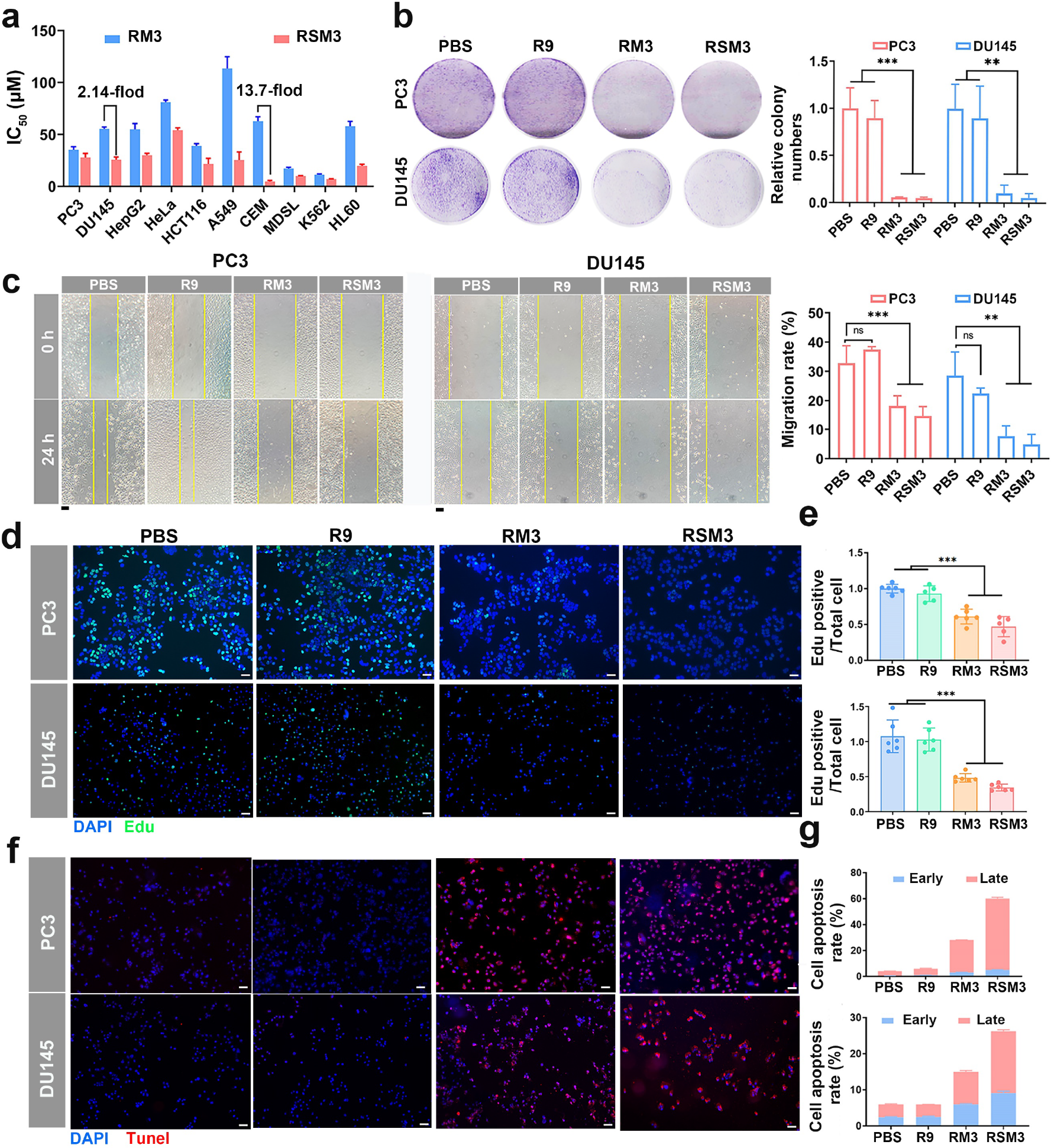
Evaluation of cytotoxic effects by METTL3-METTL14-targeting peptides. **(a)** MTT assays evaluating cell viability. **(b)** Colony formation assays and **(c)** cell migration assays in PCa cell lines after treatment with 15 μM peptide (n=3). **(d)** Representative images of EdU labeling to assess cell proliferation after treatment with 15 μM peptide. **(e)** Quantification of EdU staining. **(f)** TUNEL staining and **(g)** flow cytometry analysis of apoptosis levels in peptide-treated DU145 and PC3 cells after treatment with 15 μM peptide. For a-e mean and error (±s.d.) were obtained from at least three replicates. One-way analysis of variance (ANOVA) with Tukey’s test correction was used in statistical analysis test. ^*^P < 0.05, ^**^P < 0.01, ^***^P < 0.005, and not significant by P > 0.05. Scale bar: 20 μm.

We next investigated the cytotoxic effect of peptide inhibitors. To better understand the impacts on cell proliferation, we employed 5-ethynyl-2’-deoxyuridine (EdU) labeling with confocal laser scanning microscopy (CLSM) to assess the proportion of cancer cells that entered the S phase of the cell cycle. Image analysis indicated that PCa cells in the 15 μM **RSM3** and **RM3** treatment groups exhibited significantly weaker EdU signal, ∼35.0% and ∼47.0% that of the PBS control group (**Fig. 4d&e)**, indicating that cell proliferation was markedly reduced in the presence of these peptides. Additionally, flow cytometry analysis revealed that ∼29.2% of PC3 cells and16.9% of DU145 treated with **RSM3** were arrested in the sub-G1 phase, compared to ∼8.5% and ∼3.6% for PC3 and DU145 cells, respectively, in the PBS group (**Fig. S9**).

These findings suggested that **RSM3** could effectively induce cell cycle arrest. Subsequently, Annexin V-FITC and Terminal deoxynucleotidyl transferase dUTP Nick-End Labeling (TUNEL) staining was performed to evaluate apoptosis levels associated with 15 μM **RSM3** or **RM3** treatment. TUNEL assays clearly showed that **RSM3** could induce higher levels of apoptosis than **RM3**, although apoptosis levels in the 15 μM **RM3** treatment group were also significantly higher than in controls (**Fig. 4f**). Quantification by flow cytometry indicated that ∼60% of PC3 cells and ∼30% of DU145 entered apoptosis in the **RSM3** group, compared to 4% and ∼5% of PBS-treated PC3 and DU145 cells, respectively (**Fig. 4g**). Furthermore, a larger proportion of apoptotic cells were in the late stage, accounting for ∼90% of the total apoptotic PC3 cells. The proportion of cells with apoptosis induced by **RSM3** was approximately 2-fold higher than that of **RM3**, further supporting the higher toxicity of the stapled peptide towards cancer cells. Western blot (WB) assays confirmed that cleaved caspase-3 accumulated in peptide-treated cells (Fig. S10), further validating the apoptotic effect. Collectively, these results illustrated the anticancer activity of METTL3 peptide inhibitor by inducing cell apoptosis and cell cycle arrest.

### RSM3 interaction with METTL3-METTL14 decrease complex stability and function

To explore the interactions between the peptide inhibitor and METTL3, we used a FITC-labeled fluorescent **RM3** to track its intracellular localization. CLSM image also analysis illustrated the obvious cellular uptake of **RM3** within 2 h incubation, in contrast with the minimal cellular entry observed with the M3 control peptide lacking CPP R9 (**Fig. 5a**). This result could at least partially explain the markedly lower inhibitory effects of **M3** and **SM3** towards cancer cells compared to that of **M3** or even **SM3** in combination with the R9 cell penetration peptide. Immunostaining for METTL3 confirmed s that fluorescent signal from **RSM3** colocalized with that of METTL3 (**Fig. 5b**), while containing for METTL3 and METTL14 revealed that both proteins were obviously localized to the perinuclear region at 6 h of **RSM3** treatment^**[27]**^ **(Fig. 5c)**. Furthermore, METTL3 and METTL14 signal intensity were both decreased, suggesting that treatment with **RSM3** could potentially result in altered function and possible degradation of the METTL3-METTL14 complex. WB analysis further confirmed that METTL3 and METTL14 levels were reduced in PC3 cells at 12 h of **RSM3** treatment (**Fig. 5d**), supporting that **RSM3** could both inhibit catalytic activity and also trigger degradation of the METTL3-METTL14 complex. Subsequent RNA blot assays with m6A ELISA demonstrated that m6A modifications were significantly reduced in PC3 or DU145 cells treated with **RSM3** or **RM3**, reaching levels approximately ∼20% that detected in the R9 and PBS treatment groups (**Fig. 5e&f**). These results highlighted the potent effects of inhibiting METTL3 methyltransferase activity with stapled **RSM3** peptides.

**Figure 5.**
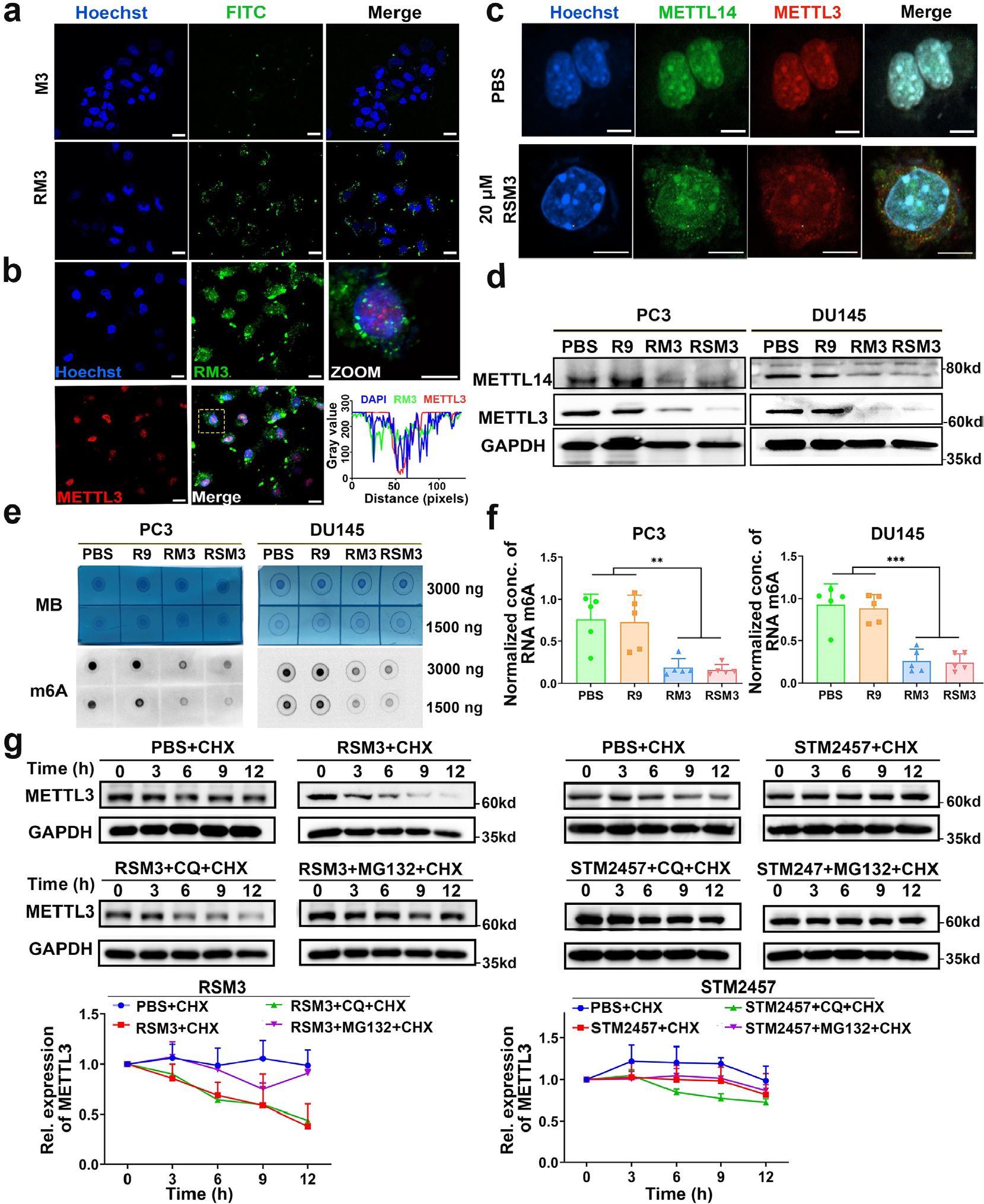
Peptide inhibitors modulate METTL3 stability and function. **(a)** Representative confocal microscopy images of PC3 cells treated with 10 μM FITC-labeled **M3** or **RM3** showing differences in cellular uptake in the presence or absence of R9 CPP. **(b)** Immunostaining for METTL3 to detect co-localization with FITC-labeled **RM3. (c)** Immunostaining for METTL3 and METTL14 in PC3 cells treated with **RSM3** and PBS vehicle control. **(d)** WB analysis to detect METTL3 or METTL14 in PC3 cells treated 25 μM **RSM3** over 12 h. **(e)** RNA blot analysis demonstrating the impact of peptide inhibitor on m6A methylation level in PC3 and DU145 cells treated with 25 μM of **RSM3** over 12 h, Methylene blue (MB). **(f)** ELISA analysis of m6A modification levels in PC3 and DU145 cells treated with **RM3, RSM3**, R9, or PBS controls. **(g)** WB analysis demonstrating the degradation of METTL3 mediated through proteasome pathway upon the treatment with CHX or MG132 and 25 μM **RSM3** over 12 h. For f,g mean and error (±s.d.) were obtained from at least three replicates. One-way analysis of variance (ANOVA) with Tukey’s test correction was used in statistical analysis test. ^*^P < 0.05, ^**^P < 0.01, ^***^P < 0.005, and not significant by P > 0.05. Scale bar 10 μm.

We next investigated the impact of peptide inhibitors on METTL3 half-life by pre-treating PC3 cells with cycloheximide (CHX) to block protein synthesis prior to **RSM3** or **RM3** administration. WB showed that **RSM3** or **RM3** treatment resulted in dramatically reduced half-life for METTL3 (**Fig. 5g**). Subsequent examination of the **RSM3**-associated degradation pathway through treatment with autophagy inhibitor (chloroquine, CQ) or proteasome inhibitor (MG132) revealed that exposure to MG132, but not CQ, could reverse the depletion of METTL3 in the **RSM3**-treated cells. By contrast, treatment with the METTL3 inhibitor, **STM2457**, had no influence on METTL3 levels in WB analysis. These results indicated that **RSM3** treatment could activate METTL3 degradation through the proteasome pathway, different from the inhibitory activity of **STM2457** on METTL3 methyltransferase activity. Previous work have demonstrated that disrupting the METTL3/METTL14 complex decreases stability, triggering ubiquitination and subsequent degradation by STUB1^[28]^. Thus, we hypothesized that the peptide might interfere with the METTL3/METTL14 complex, ultimately leading to its disruption. To investigate this, we employed a Co-Immunoprecipitation (CO-IP) assay to assess any reduction in the binding between METTL3 and METTL14 (**Fig.S11 a & b**). The results showed that the interaction of METTL3 and METTL14 was diminished with **RSM3** treatment, with quantitative analysis revealing approximately a fifty percent reduction. These findings suggest that **RSM3** effectively disrupts and destabilizes the METTL3/METTL14 complex, subsequently promoting degradation through the proteasome pathway. Taken together, these results indicated that interference and disrupting in METTL3-METTL14 interaction by **RSM3** or **RM3** could block m6A modification while also inducing the proteasomal clearance of METTL3 in PCa cells, and further suggested that the. stapled peptide **RSM3** might be an effective candidate drug for cancer therapy.

### Transcriptomic response to peptide treatment in PCa cells

To further uncover the molecular basis underpinning the effects of **RSM3** on cancer cell viability, we next performed bulk RNA-seq in PC3 cells treated with vehicle or 25 μM **RSM3** for 12 h (**Fig. 6a**). Interestingly, we identified only 89 differentially expressed genes (DEGs; |FC≥2| and FDR = 0.05) in the **RSM3** *vs*. control group comparison, indicating a highly specific transcriptomic response to m6A pathway disruption. At a less stringent threshold of FC≥1.5, 302 DEGs were detected, including 202 upregulated and 100 downregulated (**Fig. 6a**), which could be reasonably interpreted as a result of blocking m6A function in promoting mRNA decay ^[29]^. The Gene Ontology (GO) analysis of these 302 DEGs by Metascape implied that the upregulated genes were enriched in pathways involved in programmed cell death, p53 tumor suppressor signaling, differentiation, and immunity (**Fig. 6b**), whereas down-regulated genes were mainly enriched in pathways involved in promoting cancer aggressiveness, stemness, and proliferation (**Fig. 6c**). These findings were generally consistent with our recent study that oncogenic function of the m6A pathway in PCa was associated with features of tumor immune microenvironment (*e*.*g*. cytokine signaling)^[30]^. Gene set enrichment analysis (GSEA) further confirmed the GO results, showing that expression of apoptosis, inflammatory response, and p53 signaling signature genes was enriched in **RSM3**-treated cancer cells (**Fig. 6d**). In addition, the androgen receptor (AR) pathway was significantly enriched in AR^-^ PC3 cells after **RSM3** treatment (**Fig. 6e**), indicating that METTL3 inhibition by **RSM3** might be accompanied by restoration of aggressive AR^-^ cells to a more indolent AR^+^ cell state. Real-time qPCR analysis was used to validate the results of RNA-seq analysis (**Fig. 6f**), which showed upregulation of NF-κB responsive genes related to inflammation and apoptosis and downregulation of genes related to cell cycle and proliferation. In particular, the expression of luminal differentiation genes (i.e., AR, PSA, NKX3.1, CK18) was increased while expression of stemness genes decreased, further supporting the molecular differentiation phenotype associated with **RSM3** treatment **(Fig. 6f**). To verify that the effects of **RSM3** treatment were due to inhibition of m6A signaling, we checked for overlap between DEGs detected by this RNA-seq analysis and either human m6A-targeted genes recorded in the M6A2Target database ^[31]^ (**Fig. 6g**) or METTL3-bound proteins (**Fig. 6h**) identified by RIP-seq in our parallel study^[22]^. This analysis showed that DEGs detected in **RSM3**-treated PCa cells were indeed significantly enriched with genes subject to m6A modulation, thus illustrating the on-target effects of METTL3 inhibition by **RSM3**. Furthermore, we also noted that MAPK signal cascade genes were dramatically overrepresented among the upregulated GO terms in **RSM3**-treated cells (**Fig. 6b**), implying that **RSM3** may deactivate MAPK activity. Western blot analysis further showed lower phosphorylation levels of p42/44 in the **RSM3** group (**Fig. 6i**), indicating that **RSM3** treatment resulted in decreased activity of ERK/MAPK. Collectively, our data supported that METTL3 inhibition by **RSM3** could retard cancer cell growth and evoked apoptosis via inhibition of MAPK mitotic pathway, together with induction of differentiation, in an m6A-dependent manner.

**Figure 6.**
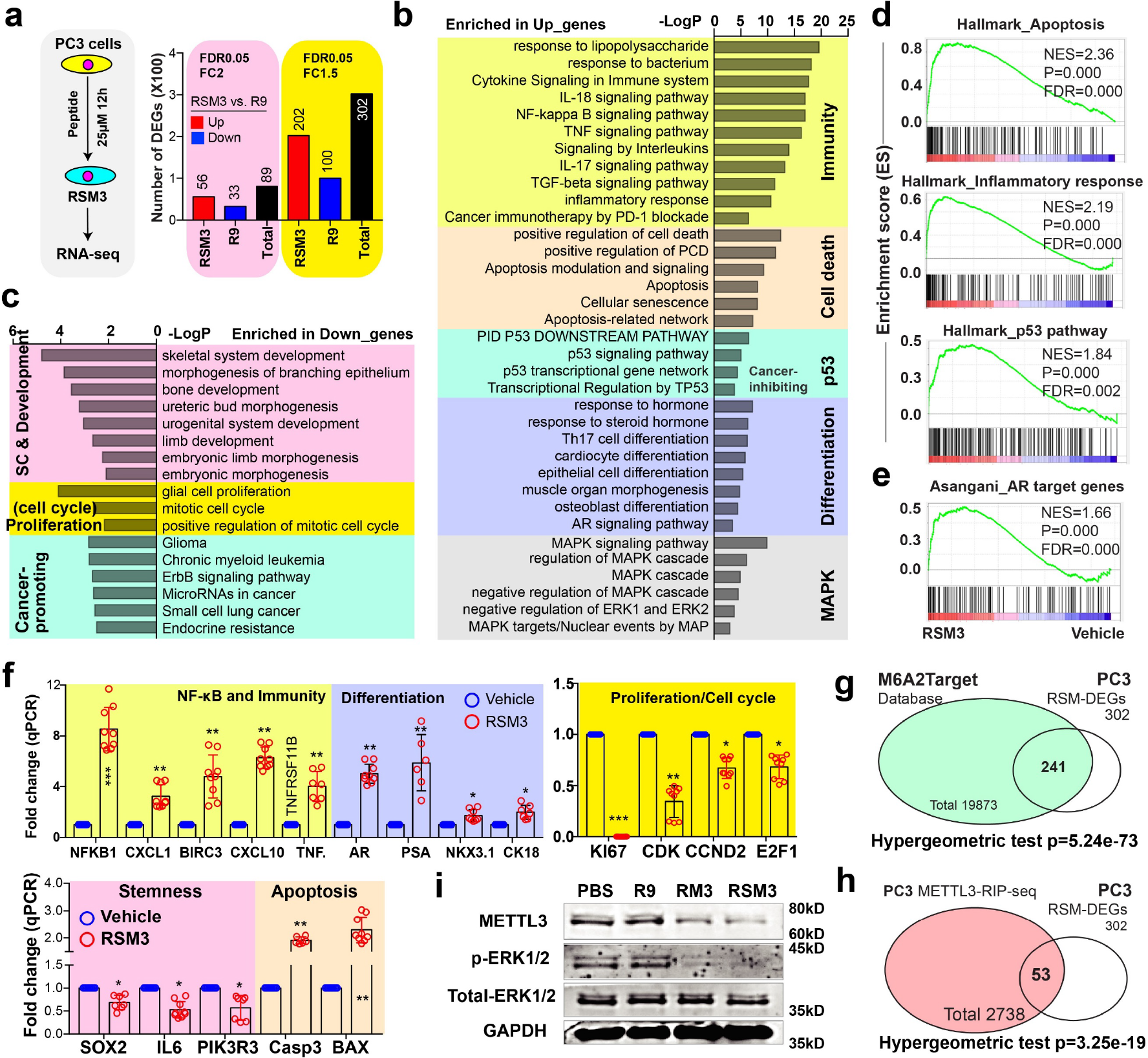
Molecular impact of METTL3-targeting peptide on cancer cell transcriptome. **(a)** Effect of **RSM3** on PCa transcriptome *in vitro*. Shown are schema of RNA-seq experiments (left) and total DEGs (right) identified by two different but stringent statistical thresholds. GO analysis of upregulated **(b)** and downregulated **(c)** DEGs in PC3 cells treated with or without 25 μM **RSM3**. Terms with p-value <0.01, minimum count 3, and enrichment factor >1.5 (the ratio between observed count and the count expected by chance) were considered significant. The terms with similar descriptions and functions were grouped into different functional categories. **(d) and (e)** GSEA showing enrichment of the indicated gene signature in **RSM3**-*vs*. vehicle-treated PC3 cells. **(f)** qPCR analysis of indicated genes in PC3 cells treated with or without 25 μM **RSM3**. Overlap between 302 DEGs and human m6A-targeted genes in M6A2Target database (**g**) and METTL3-bound genes identified in PC3 cells (**h**). Statistics was calculated by hypergeometric test. **(I)** WB analysis of p42/44 MAPK phosphorylation in PC3 cells treated with vehicle or peptide inhibitors at 25 μM for 12 h. For **f** mean and error (±s.d.) were obtained from at least three replicates. One-way analysis of variance (ANOVA) with Tukey’s test correction was used in statistical analysis test. ^*^P < 0.05, ^**^P < 0.01, ^***^P < 0.005, and not significant by P > 0.05.

### RSM3 safety and anti-tumour efficacy *in vivo*

To assess safety, we examined its acute toxicity *in vivo* through a single intraperitoneal injection of **RSM3** (50 mg/kg) into C57/BL mice. After 24 h, routine blood analysis showed no significant differences in white blood cells (WBC), platelet (PLT), or red blood cell (RBC) levels between mice injected with **RSM3** and those injected with **RM3, R9**, or PBS. Although hemoglobin (HGB) levels were significantly lower in the **RSM3** group compared to the PBS control group, HGB values remained within acceptable limits (**Fig. 7a**). Histopathological examination of major organs, including the heart, liver, spleen, lung, and kidney, from mice treated with the peptide inhibitors, showed no significant abnormalities or lesions in any treatment groups (**Fig. S12**). Multiple-dose injections of the peptide were also conducted to thoroughly assess *in vivo* toxicity. The results from blood routine analysis and histopathological examination of major organs revealed no significant signs of toxicity (**Fig. S13**). These results strongly suggested that the peptide inhibitors conferred no obvious toxicity to mice.

**Figure 7.**
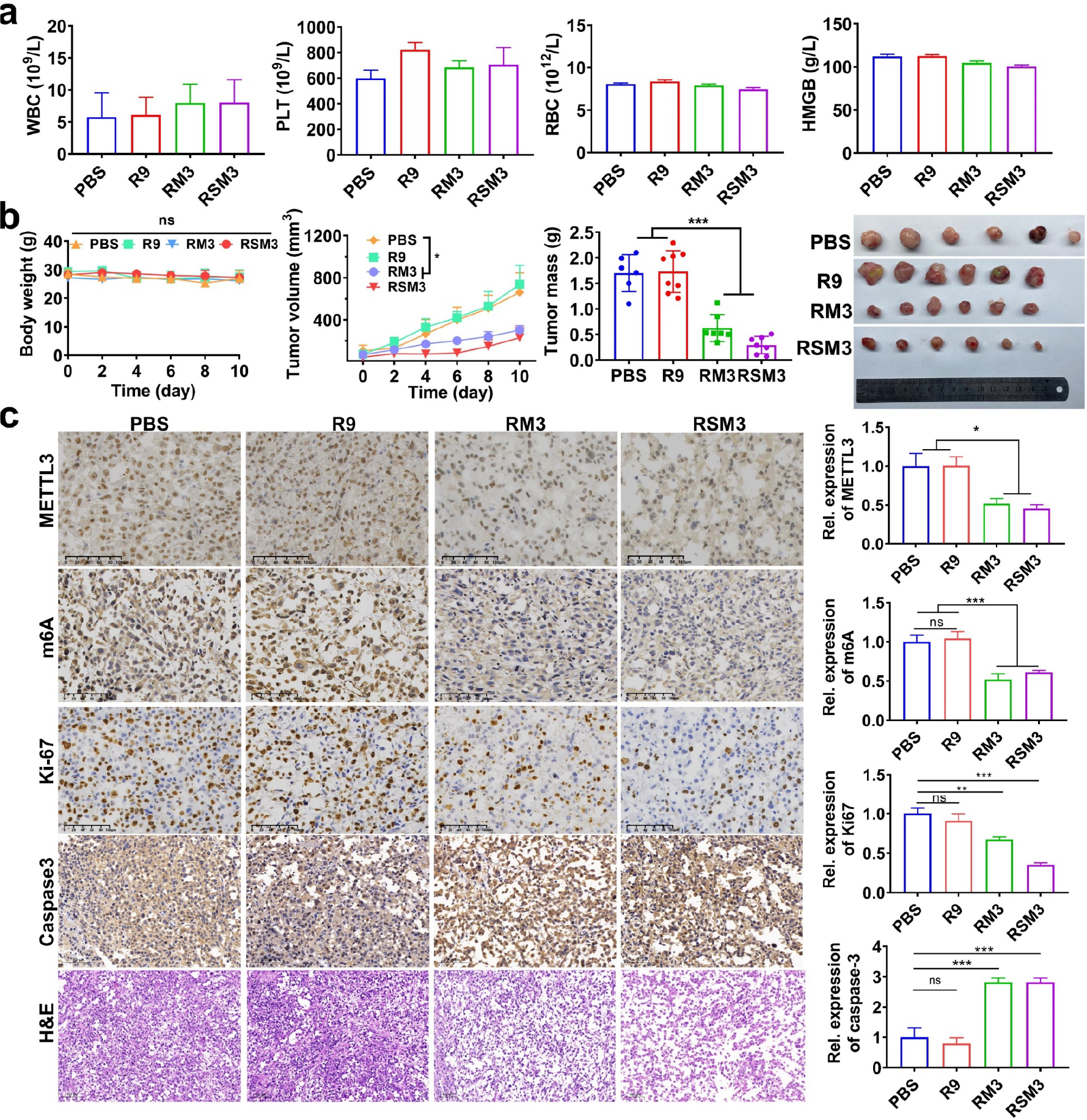
Toxicity and anti-tumor efficacy studies of peptide inhibitors. **(a)** Blood routine analysis (WBC, PLT, RBC, HMGB) of C57/BL mice after a single administration of 50 mg/kg **RSM3** to assess the toxicity of the peptide inhibitor. **(b)** The change in body weight and tumor index (including tumor volume and tumor mass) was recorded and analyzed during the therapeutic process to demonstrate the tumor-suppressive effects of the peptide inhibitor in the PC3 cells xenograft tumor model. **(c)** Immunohistochemistry (IHC) image and quantification of expression analysis of METTL3, Ki67, caspases-3 and m6A in PC3 cells xenograft mice tumor tissue after peritumoral injecting with peptide inhibitor (20 mg/kg) for 10 days, scale bar:100 μm. For **a-c** mean and error (±s.d.) were obtained from at least three replicates. One-way analysis of variance (ANOVA) with Tukey’s test correction was used in statistical analysis test. ^*^P < 0.05, ^**^P < 0.01, ^***^P < 0.005, and not significant by P > 0.05

We then evaluated the anti-cancer activity of **RSM3** in PC3 xenograft tumor model. Tumor-bearing mice were given a single peri-tumoral injection of 100 μL **RM3, RSM3** (20 mg/kg), or the R9 or PBS control components, every two days for two weeks. Measurements of tumor size and mouse body weight throughout the 2-week experimental period indicated that **RSM3** or **RM3** treatment resulted in significant suppression of tumor growth compared to the R9 or PBS control groups (**Fig. 7b)**. More specifically, tumor volumes were 2∼3 fold smaller in the **RSM3** group compared to controls, with the stapled **RSM3** peptide exhibiting higher efficacy than the non-stapled **RM3** peptide, suggesting that increased stability through stapling could enhance the peptide inhibitory activity *in vivo*. Throughout the therapeutic course, the body weights were unaffected in all groups, and histopathological examination of organs (heart, liver, spleen, lung, kidney) found no significant abnormalities or lesions, indicating that none of the compounds had noticeable toxicity in the tumor-bearing mice (**Fig. S14**). Immunohistochemical analysis of tumor samples showed that treatment with either **RM3** or **RSM3** resulted in a ∼50% reduction in METTL3 expression and m6A levels compared to the R9 and PBS groups. Additionally, caspase-3 expression increased by ∼1.8-fold, while Ki67 decreased by ∼65%, indicating that tumor growth was likely suppressed in the **RSM3** or **RM3** groups via induction of apoptosis and inhibition of proliferation (**Fig. 7c**). Furthermore, using the ASPC-1 cell line and ASPC-1 xenograft tumor model of pancreatic cancer to evaluate generalizability of peptide inhibitors, we found that **RSM3** also exhibited substantial anti-tumor activity towards ASPC-1 cells (**Fig. S15**, IC_50_ = 33.8 ± 7.0 μM) and ASPC-1 xenograft tumors (**Fig. S16)**, with significantly lower tumor volumes, and ∼50% lighter tumor weights in the **RSM3**-treated group compared to the PBS- and R9-treated control groups. We speculated thatthese observations might be due to a commorole of METTL3 in promoting pancreatic cancer development and metastases^[32]^.

Based on our above findings, we next sought to compare the anti-tumor effects of **RSM3** with that of the previously reported small molecule inhibitor of METTL3, **STM2457** ^[14]^. Both **STM2457** and **RSM3** exhibited potent toxicity towards PC3 and DU145 cells (**Fig. S17**), and **RSM3** displayed higher toxicity, with lower IC_50_ values, than the **STM2457** in PC3 (26.5 ± 1.2 *vs* 95.5 ± 3.6, **RSM3** *vs* **STM2457**) and DU145 (33.6 ± 1.5 *vs* 149.3 ± 10.1, **RSM3** *vs* **STM2457**) cells. Subsequent RNA blot assays revealed a significant reduction in m6A levels in PC3 cells treated with either **RSM3** or **STM2457**. Notably, **RSM3** exhibited a marginally superior m6A inhibition effect compared to **STM2457** (**Fig. S18**). To this end, **STM2457** or **RSM3** were administered to PC3 xenograft model mice at the same dosage (20 mg/kg), which demonstrated that both **STM2457** and **RSM3** could effectively suppress tumor growth in PCa tumors *in vivo* (**Fig. S19)**. However, tumor mass was significantly lower in the **RSM3** group compared to **STM2457**-treated tumors (**Fig. S19c**), that is, average tumor mass in the **RSM3** group was ∼37% that in the PBS group, while average tumor mass in the **STM2457** group was 70% that of controls. These results demonstrated that peptide inhibitors of METTL3 can effectively suppress prostate and pancreatic tumor development *in vivo*. Moreover, these findings in xenograft tumor model supporting the potent disruption of METTL3 activity and stability by **RSM3** suggest that these peptide inhibitors function through a different and potentially more reliable pathway than small molecule METTL3 inhibitors.

## Conclusion

Our current study focused on the development of a Bph-linked stapled peptide inhibitor targeting the m6A methyltransferase, METTL3, in cancer cells. Comprehensive physicochemical and functional characterization in multiple cancer cell lines *in vitro* demonstrated that the **RSM3** stapled peptide displayed relatively high affinity towards the METTL3 binding site for complex formation with METTL14, reasonably high stability in denaturing conditions compared to other candidates, and significant cytotoxicity and anti-tumor activity *in vitro* and *in vivo*. Notably, **RSM3** binding with METTL3 not only disrupted its activity, but also activated its proteasomal degradation, resulting in substantially reduced mRNA m6A global levels, *in vitro* and *in vivo*. In treatments of two different PCa cell lines, **RSM3** exhibited potent inhibition of cell proliferation and migration, supporting its potential as a therapeutic agent. Transcriptomic analysis indicated that **RSM3** could trigger inflammatory response pathways, activating p53 signaling, ultimately inducing cell cycle arrest and ultimately programmed cell death via apoptosis. We also validated that **RSM3** deactivates MAPK signaling. In PC3 Xenograft Model mice, treatment with **RSM3** twice a day of 20mg/kg could reduce METTL3 expression and promote apoptosis in the tumor microenvironment *in vivo*, thus providing compelling evidence if its efficacy. Furthermore, **RSM3** could provide significantly higher cytotoxicity and anti-tumor activity towards PCa *in vivo* and *in vitro* compared to the small molecule METTL3 inhibitor, **STM2457**, potentially due to differences in the mechanism of action. The combined properties of high METTL3 inactivation and prolonged stability in the tumor cells collectively support the potential for broad application and generalizability of **RSM3** in treating a range of METTL3-dependent cancers ^[12a]^. In addition, the innovative design strategy for **RSM3** provides a theoretical framework for developing future research tools and targeted therapies, including other diseases linked to dysregulation of m6A methylation, with broader implications for targeting other complexes with oncogenic function in future therapeutic development.

## Supporting Information

Materials and methods, synthesis, and characterization of all peptides.

## Acknowledgements

This work was supported by the National Natural Science Foun-dation of China (21975068 to J.S. and 81972418 to D.Z.), the National Youth Talent Support Program (202309460011), Natural Science Foundation of Hunan (2020RC3017 and 2022JJ10008 to J.S., 2021JJ10028 to D.Z.), Shenzhen Science and Technology Program (JCYJ20210324122403010 to J.S., JCYJ20220530160410024 to D.Z.).We would like to thank the Analytical Instrumentation Center of Hunan University for Assistance in Confocal Microscopy, and we also very grateful to Ping Yin’s group for providing METTL3-METTL14 protein.

## Conflict of interest

J.S, D.Z, L.Z, Y.F, M.W, and H.H are inventors on pending patent applications (No. 202211104956.7, and 202211105358.1), the authors declare no other competing financial interest.

## Notes

### Summary of Updates

1. We have amended textual and visual inaccuracies, consequently enhancing the overall quality of the manuscript. 2. We elucidate the conceptual framework of peptide design and the construction process of stapled peptides. 3. We incorporated co-ip and biotinylated peptides pull down assay to study work mechanism of peptide.

