## Supplemental information for "A Stapled Peptide Inhibitor of METTL3-METTL14 for Cancer Therapy"

# of

### Table of Contents


**Materials and Experimental Methods**

**Table S1.** $IC_{50}$ value of peptide inhibitors towards multiple cancer cell lines.
**Figure S1-3.** HPLC and mass spectra of peptides studied.
**Figure S4.** The principle of stapled peptide design..
**Figure S5.** Synthetic routes of stapled peptides.
**Figure S6.** Evaluation of **RM3** and **RSM3** binding to METTL3/METTL14 by Octet® BLI Systems.
**Figure S7.** Biotinylated peptide pull-down assay demonstrating RM3 binding to the METTL3/14 complex in cell lysate.
**Figure S8.** The MTT assay determinate the cytotoxicity of peptide inhibitors towards multiple cancer cell lines.
**Figure S9.** Flow cytometry investigating the effect of peptide inhibitors on the cell cycle.
**Figure S10.** WB analysis detecting the activation of caspase3 in PC3 after peptide inhibitors treatment.
**Figure S11.** Co-IP assessing the interaction between METTL3 and METTL14 in the absence or presence of **RSM3**.
**Figure S12.** H&E staining of organ sections after once dose injection with peptide for safety evolution.
**Figure S13.** Blood routine and H&E staining after multiple dose injection with peptide for safety evolution.
**Figure S14.** H&E staining of organ sections from PC3-xenograft mice after peptide administration.
**Figure S15.** The MTT assay determinate the cytotoxicity of peptide inhibitors towards pancreatic cancer cell ASPC-1.
**Figure S16.** Peptide inhibitor suppresses tumor growth in pancreatic cancer cell ASPC-1 xenograft tumor model.
**Figure S17.** Cytotoxicity of **RSM3** compared to small molecular inhibitor **STM2457** in a prostate cancer model.
**Figure S18.** RNA blot assay showing the different impact of **RSM3** and **STM2457** on m6A level in PC3 cells.
**Figure S19.** Therapeutic effect of **RSM3** compared to small molecular inhibitor **STM2457** in a prostate cancer model.

#### Materials and Experimental Methods

##### Materials

Rink Amide (AM) Resin, Fmoc-protected amino acids and O-Benzotriazole-N, N, N', N'-tetramethyl-uronium-hexafluoro-phosphate (HBTU) were purchased from Hitch Biotech Shanghai Co., Ltd. Trifluoroacetic acid (TFA), acetonitrile, methanol, hexane, piperidine, and purified water were purchased from inno-chem. The pull-down assay was conducted using streptavidin-labeled magnetic beads (biolinkedin, L-1012). All the solvents and reagents were used directly as received from commercial sources without further purification

##### Peptide Synthesis

All peptides were synthesized by standard Fmoc solid-phase peptide synthesis on a CSBio 136S peptide synthesizer, with Rink Amide AM Resin and activation by HCTU. After the synthesis of the peptide chain, the resin should be washed successively with DMF, DCM, MEOH and n-hexane, Finally, the peptide chain without side chain protecting groups was cleaved from the resin with TFA/thioanisole/H2O (95:2.5:2.5) for 3 h under nitrogen atmosphere. Crude peptide was obtained by concentrating the filtrate and precipitating it with cold ether. The crude product was purified by reversed phase high performance liquid chromatography (RP-HPLC) and lyophilized to obtain dry powder. All purified peptides were analyzed using analytical HPLC and MALDI-TOF MS.

Following a similarly procedure described above, fluorescently labeled peptides were linked to the N-terminal of the peptide on the resin with Fluorescein-succinimidyl ester (NHS-FITC) (1,5 eq.) and DIEA (4 eq.) in DMF for overnight with constant shaking at room temperature. All purified peptides were analyzed using analytical HPLC and MALDI-TOF MS.

##### Synthesis of SM3 and RSM3

Cross-linking reactions were performed by incubating M3 and RM3 peptides with 1.5 equivalents of P-bromo-methyl biphenyl (bhp) in a mixture of acetonitrile and 50 mM $NH_4HCO_3$ (1:1), pH 8.5, at a final peptide concentration of 1.0 mM. The reaction mixture was then incubated at room temperature for 2.0 h. The progress of the reaction was monitored using Analytical HPLC. Once the reaction was complete, the solvents were evaporated and any excess cross-linkers were removed by washing with diethyl ether. The cross-linked peptides were subsequently purified using preparative HPLC.

**Molecular Dynamics Simulation**

The initial model for simulations was constructed from the docking structure. The Amber14SB force field (1) was utilized for molecular dynamics simulations, counterions (Na+, Cl-) were added to neutralize net system charge, before solvating in a TIP3P (2) water box with ⩾10 Å buffer from the protein surface. Prior to production, the system underwent energy minimization using steepest descent and conjugate gradient algorithms (5,000 cycles) (3,4), followed by gradual temperature annealing from 10 to 300 K (0.05 ns) under weak harmonic restraints (15 kcal/mol/Å2). Density and pressure equilibration were subsequently performed for 1 ns under isothermal-isobaric conditions (300 K, 1 atm) using Langevin dynamics (5) and Berendsen/Parrinello-Rahman barostats (6,7). After verifying adequate equilibration, 200 ns production runs were started using GROMACS 2022.4 (8). GMX_MMPBSA was subsequently utilized to compute residue-wise free energy decomposition over the aggregated trajectories, elucidating key binding determinants (9).

**Circular Dichroism (CD)**

CD spectra were obtained using a Jasco J-1500 spectrometer. Spectra were collected for all peptides in a 1 mm quartz cell with wavelength. The 150 μM peptides were prepared in D.I. water and chilled on ice for at least 0.5 h. The mean residue ellipticity, [θ], was calculated using the following formula: $[\theta]=(\theta_{obs}/10lc)/r$, where $\theta_{obs}$ is the observed ellipticity in millidegrees, l is the length of the cell in centimeters, c is the concentration in molarity, and r is the number of residues.

**Microscale Thermophoresis (MST) Assay**

All measurements were conducted using the Monolith NT 115 instrument. Each measurement utilized a total volume of 10 μL loaded into standard capillaries. The ligand peptide was prepared by diluting it into a series of concentrations from a 5 mM stock solution. Prior to the measurements, the purified METTL3-METTL14 protein was diluted in assay buffer and labeled using the MO RED-NHS Protein Labeling Kit. After a 5-minute incubation period, the peptide inhibitors were added to achieve final concentrations of 50 nM for METTL3-METTL14 and a range of concentrations for the peptide inhibitor stock (ranging from 250 μM to 1.95 μM). Following an additional 10-minute incubation, the measurements were immediately taken using the same settings as previously described. The resulting data were processed using the MO Affinity Analysis software v2.3.

#### <u>Serum Stability and Hemolysis Studies</u>

##### Stability

Peptide were dissolved in 1:3 serum: buffer (20 mM Tris-HCl, 100 mM NaCl, pH 7.4) to a final concentration of 200 μM. Samples were then incubated at 37 °C in an incubator. At 0, 0.5, 1, 2, 4 or 8 h of incubation a 100 μL aliquot was removed and diluted with an equal volume of 15 wt% TCA in water, and the sample was kept on ice for 15 min. Samples were then centrifuged at 13,000 rpm for 10 min, and supernatant collected and subjected to analytical MS.

##### Hemolysis

Hemolysis investigations were conducted by procuring freshly drawn blood from healthy human volunteers into heparinized tubes, followed by subjecting it to centrifugation at a speed of 3,000 rpm for a duration of 10 minutes at a temperature of 4 °C. Red blood cells (RBCs) were subjected to three washes with a hemolysis buffer composed of 10mM Tris, 150 mM NaCl, pH 7.4. Subsequently, a solution of RBCs in the hemolysis buffer, with a concentration of 0.25% v/v, was meticulously prepared. In a 96-well plate, a volume of 75 µL of the RBC solution was admixed with an equivalent amount of a 2x peptide solution, previously dissolved in the hemolysis buffer, to initiate the assay. Negative and positive controls were established using a blank and a buffer containing 1% Triton-X100, respectively. The samples were subjected to incubation for a duration of 24 h, accompanied by gentle agitation. Subsequently, the plates were centrifuged at 4,000 rpm for 10 minutes at a temperature of 4 °C to precipitate intact RBCs. Following this, a volume of 100 μL of the supernatant from each well was extracted and transferred to an empty 96-well plate. The absorbance was then measured at a wavelength of 415 nm utilizing a microplate reader.

#### Cell Viability Assay

DU145 and PC3 cells were seeded in 96 plates at a density of 8000/well and continued to culture overnight at a 37°C, 5% CO2 incubator. Peptide stock solution was prepared in water and a series of concentrations of peptide work solution. Cells were incubated with peptide work solution for 24 h. Then the 100 μL 10 %v/v CCK8 solution was added to the plate well for 1 hour. After incubation, the plates were subjected to the UV plate reader to record 450 nm data. The absorbance of the negative controls was subtracted from each sample as a blank, and the percent viability was calculated as follows: (Absorbance peptide-treated cells / Absorbance untreated cells) ×100. GraphPad Prism 7.0 software was used to fit cytotoxicity curves and calculated $IC_{50}$ values with a non-linear regression model.

#### Laser Scanning Confocal Microscope

##### Peptide uptake

Cells were seeded on 35 mm confocal dishes at a density of 8000/well and continued to culture for 24 h in a 37 °C incubator with 5% $CO_2$. Then, the cells were washed twice with PBS buffer and incubated with 1 mL peptide Prepared-work solution (30 μM) at 37 °C with 5 % $CO_2$ for desired time. The incubated solution of cells was removed, and the following was washed twice. Then the cells were stained with 2 μg/mL Hoechst 33342 for 15 min, cell imaging was performed on an Olympus FV1200 microscopy with 60X oil objective, and the cells were maintained in a cell living imaging buffer during confocal imaging.

##### EdU Staining

The cells were cultured in 12-well plates at an appropriate density. Subsequently, the cells were treated with the 15 μM peptide inhibitor for the specified duration. To label proliferating cells, a 2 X EdU working solution (20 μM) pre-warmed to 37 °C was added to each well, resulting in a final concentration of 1 X EdU in the plate. The cells were then incubated for 2 h. After the EdU labeling period, the culture medium was removed, and 1 mL of fixative solution (PFA) was added to each well. The cells were fixed for 15 minutes at room temperature. The fixative solution was then removed, and the cells were washed three times. Finally, 0.5 mL of Click reaction solution was added to each well, and the plate was gently shaken to ensure even coverage of the sample. The images of the EdU staining were captured using the EVOSTM M5000 imaging system.

**Tunel Staining**

The cells were cultured in 12-well plates at an appropriate density. Following the treatment with the 15 μM peptide inhibitor for the designated time period, the cells were fixed using PFA and subjected to Tunel (K1134, APExBIO, Houston, USA) staining for 60 minutes at 37 ℃. DAPI staining was used to visualize the cell nuclei. The fluorescence microscope was utilized to count the number of Tunel-positive cells. The ratio of Tunel-positive cells to the total cell count was calculated to evaluate the level of cell apoptosis.

**Cell Scratch Assay**

The cells were seeded in a 12-well plate and cultured for a duration of 24 h, facilitating their growth until reaching complete confluence. Then the scratch of cells was made across the cell monolayer using a sterile pipette tip. The detached cells were removed by washing with an FBS-free medium, and a fresh medium containing the desired treatment (peptide inhibitor or control molecule) was added. The cells were then incubated under appropriate conditions for 12 h, and finally images of the scratched area were captured using a microscope. The migration of cells into the scratched area was assessed by measuring the width of the different groups. The extent of cell migration was quantified using **ImageJ** software.

**Western blotting (WB)**

Cells were seeded into a 6-well plate at a density of $1x10^5$ cells per well and cultured for

24 h. Following treatment with the 15 μM peptide or inhibitor for the appropriate duration, the treated cells were lysed using 1x passive buffer supplemented with protease inhibitors such as PMSF and a phosphatase cocktail. The concentration of total protein in the lysates was determined using Quick Start Bradford 1x Dye Reagent, and the protein concentration was calculated accordingly. Subsequently, the protein samples were denatured by heating them at 95-100°C for 5 minutes and subjected to the Western blotting assay.

**Peptide safety and Therapeutic Effect Study in Xenograft Models**

All animal research conducted in this study was approved by the Ethics Committee for Animal Experiments at HNU University (Approval No. HNU-IACUC-2021-102).

**Safety study**

Six-week-old male C57/BL mice were procured from GemPharmatech LLC. (JiangSu). In the evaluation of acute toxicity, the mice received a single injection of 50 mg/kg of the peptide. After 24 h, the mice were euthanized for the collection of organs and blood samples for subsequent analysis. The gathered data were further examined to assess the peptide safety. For the assessment of long-term toxicity, mice were injected with peptide 50 mg/kg every two days for a duration of 20 days.

**Therapeutic Effect**

Six-week-old male BALB/c nude mice were were purchased from GemPharmatech LLC. (JiangSu) for the study. PC3 cells were subcutaneously injected into the back of the mice at a density of $1x10^6$ cells in 50 μL PBS. Once the tumor volume reached approximately 100 $mm^3$, the mice were randomly divided into four groups and administered the drug once every two days at dosage of 20 mg/kg. The weight of the mice and the tumor volume were recorded every two days throughout the experiment. At the end of the study, the mice were euthanized, and tumor mass was measured. Organs and tumor tissue samples were collected for further analysis, including H&E staining and immunohistochemical analysis.

**Table S1.** $IC_{50}$ value of peptide inhibitors towards multiple cancer cell lines.

| | RM3 ($IC_{50}$, μM) | RSM3 ($IC_{50}$, μM) |
|---|---|---|
| **PC3** | 35.3±2.7 | 27.7±4.0 |
| **DU145** | 55.3±1.9 | 25.8±2.4 |
| **HepG2** | 54.9±5.6 | 29.8±2.0 |
| **HeLa** | 81.0±2.1 | 54.0±2.3 |
| **HCT116** | 38.8±2.3 | 21.6±5.3 |
| **A549** | 113.5±11.3 | 25.4±7.62 |
| **CEM** | 62.6±4.4 | 4.6±1.03 |
| **MDSL** | 17.1±1.1 | 10.0±0.35 |
| **K562** | 11.2±0.9 | 7.0±0.25 |
| **HL60** | 58.0±4.6 | 19.8±1.5 |

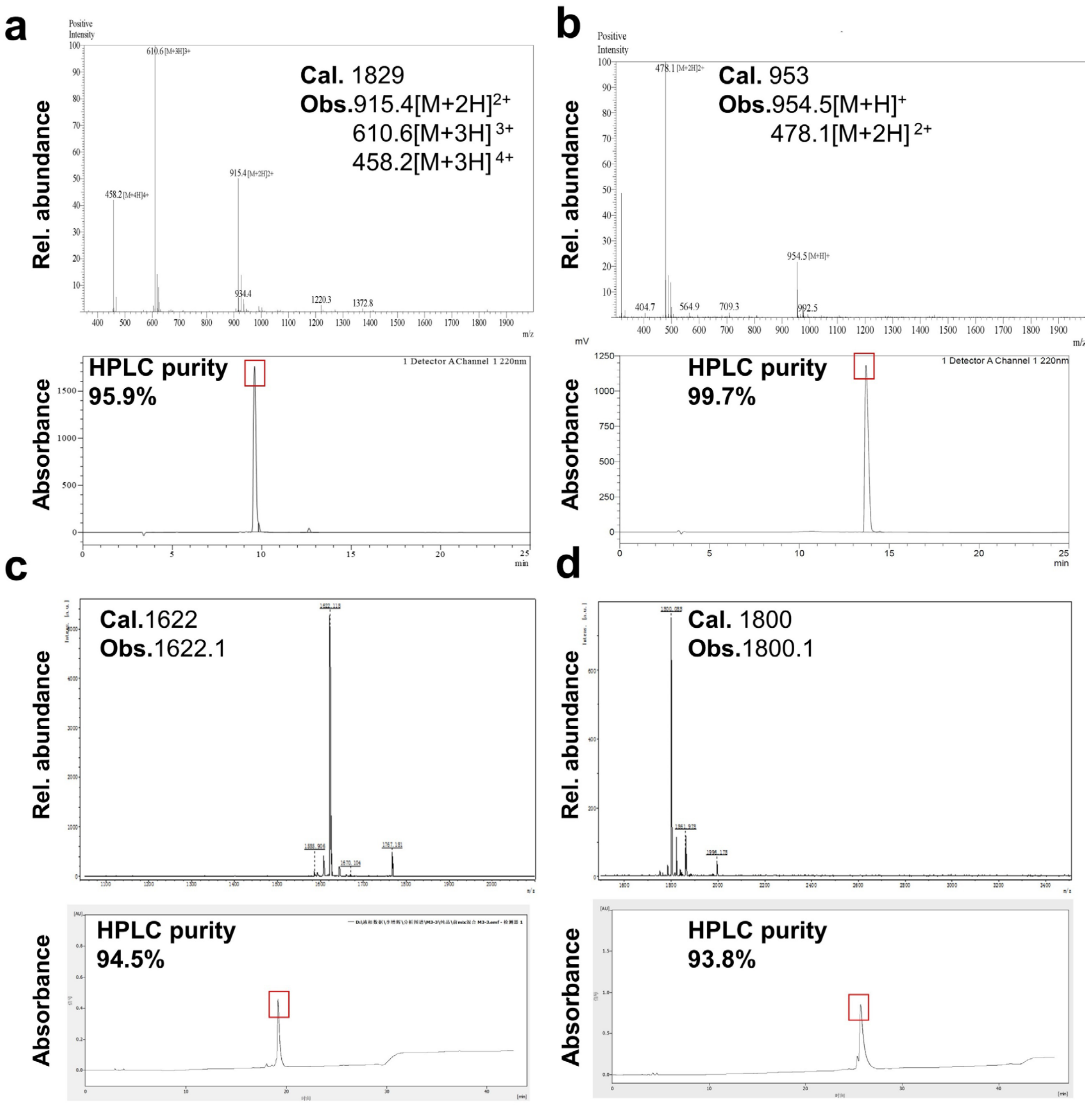


**Figure S1.** Analytic HPLC and mass spectra of peptides studied. **(a) M1**, **(b) M2**, **(c) M3**, **(d) SM3,** respective shown in this figure.

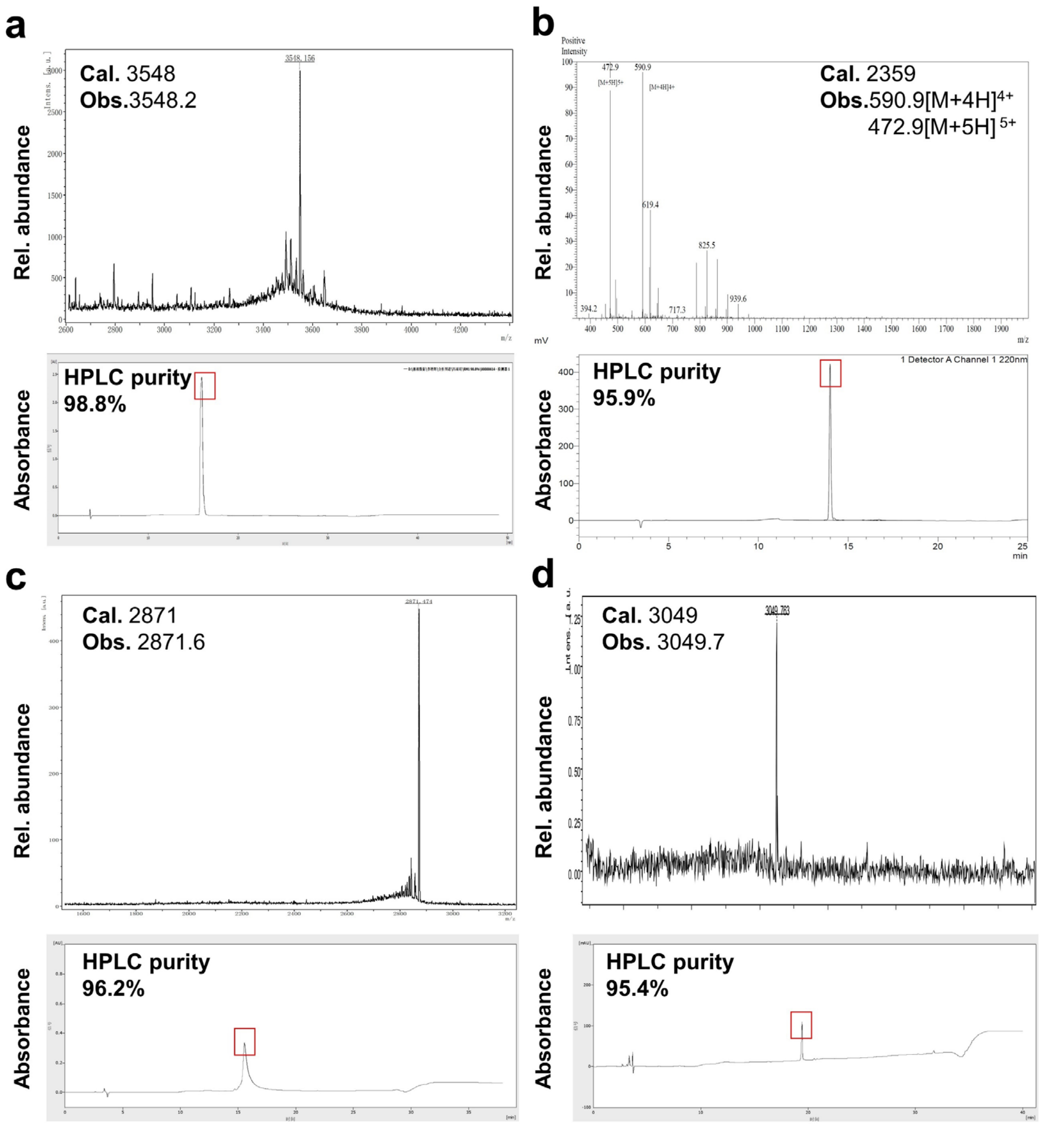


**Figure S2.** Analytic HPLC and mass spectra of peptides studied. **(a) RM1**, **(b) RM2, (c) RM3, (d) RSM3,** respective shown in this figure.

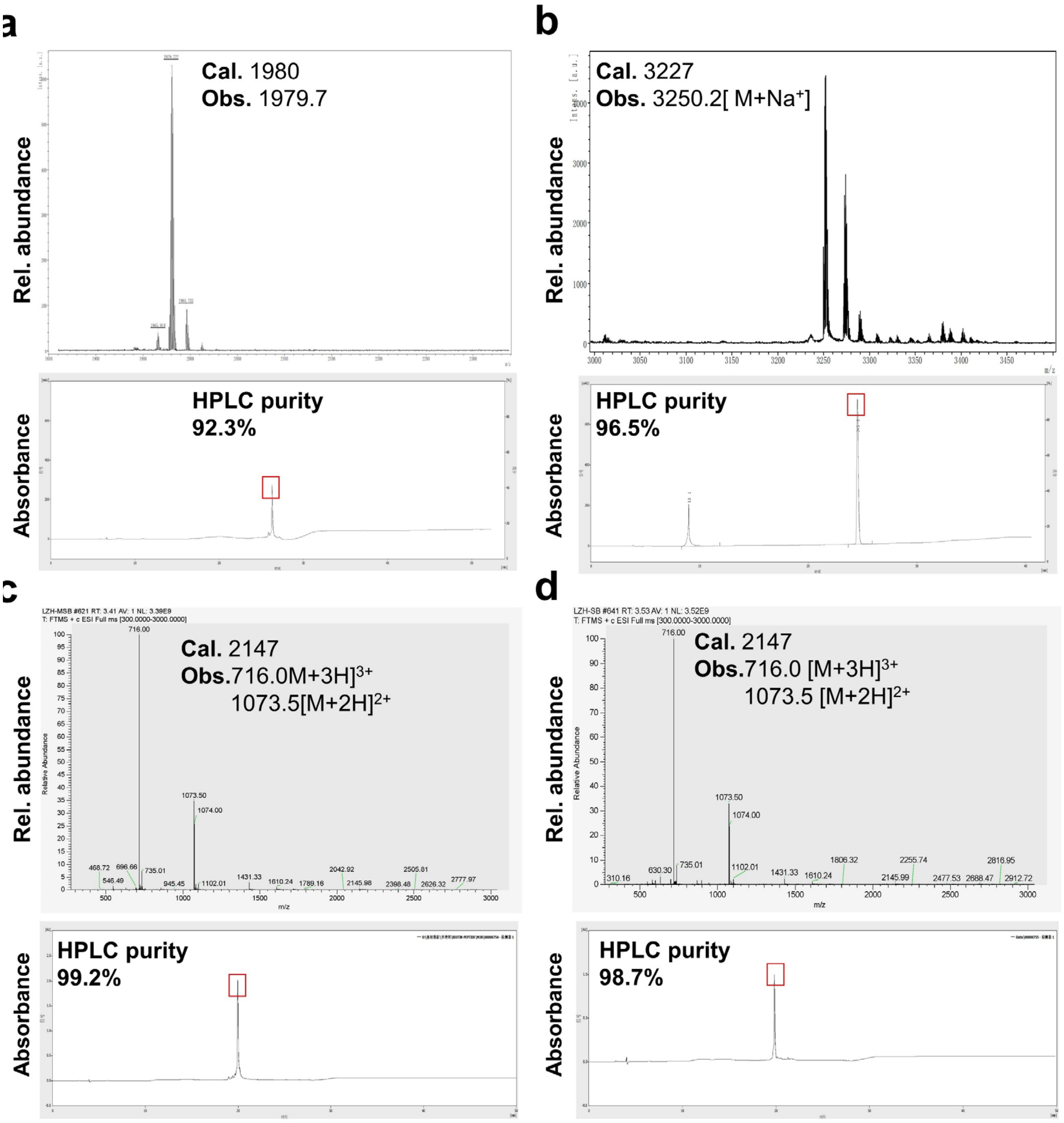


**Figure S3.** Analytic HPLC and mass spectra of peptides studied. **(a) M3-FITC, (b) RM3-FITC, (c) Bintin-M3, (d) Biotin-scrambled peptide**, respective shown in this figure.

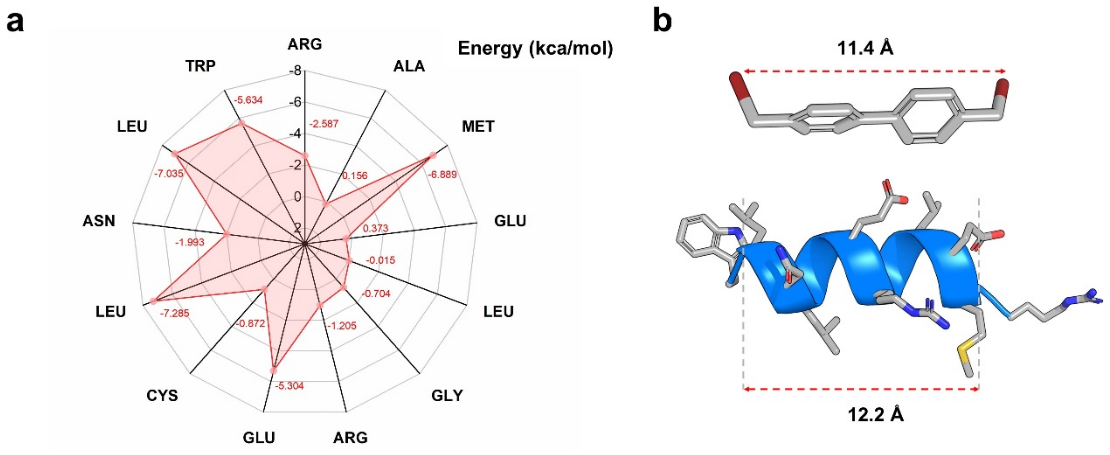


**Figure S4.** The principle of stapled peptide design. **(a)** The analysis of residues binding energy at the interface between peptide and receptor protein. **(b)** Geometric arrangement of the linker (Bph) and specific residue in space.

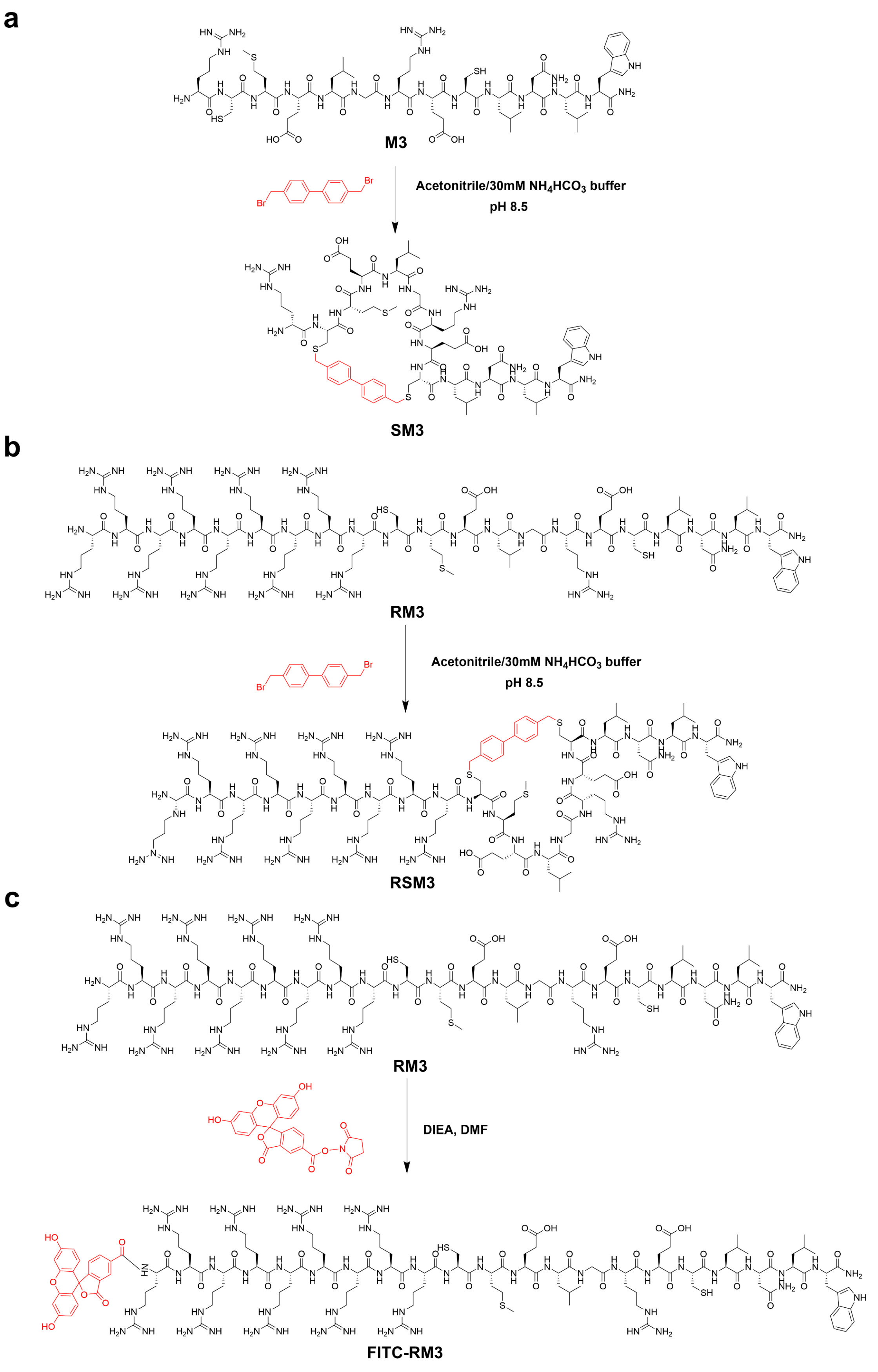


**Figure S5.** Synthetic routes of staple peptide **SM3 (a)** and **RSM3 (b)**, **(c) FITC-RM3**.

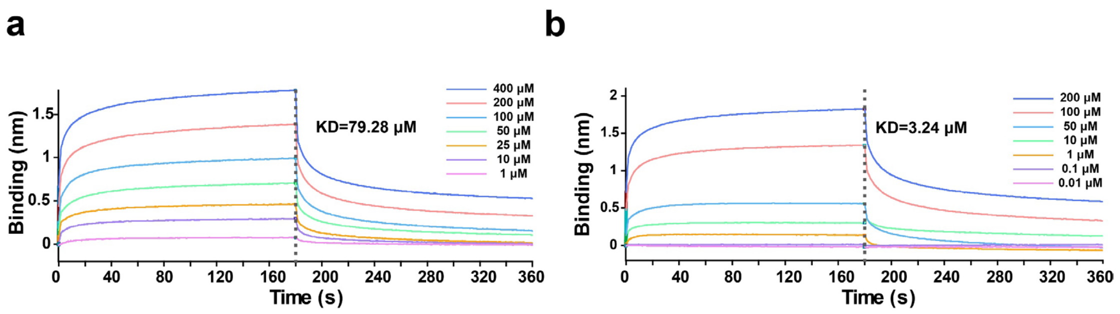


**Figure S6.** Evaluation of **(a) RM3** and **(b) RSM3** binding to METTL3/METTL14 using Octet® BLI Systems.

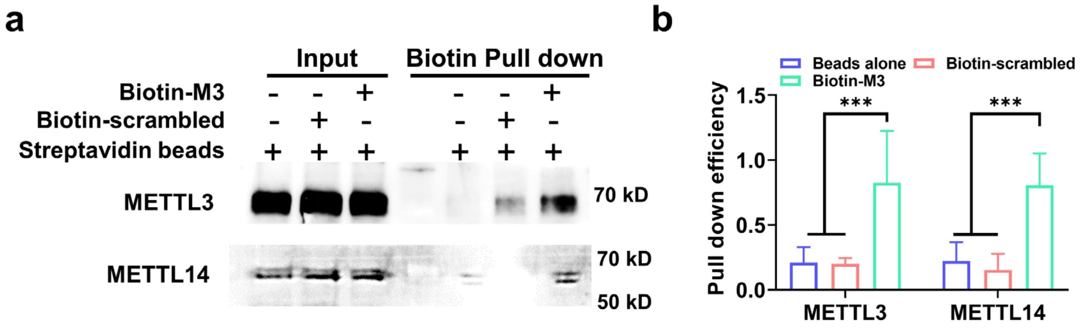


**Figure S7.** Biotinylated peptide Pull-down assay demonstrating **(a)** M3 binding to the METTL3/14 complex in cell lysate and **(b)** Quantitative analysis of pull-down efficiency. The mean and error (±s.d.) were obtained from three replicates. Statistical analysis was performed using one-way analysis of variance (ANOVA) with Tukey's test correction. *$P < 0.05$, **$P < 0.01$, ***$P < 0.005$, and not significant when $P > 0.05$.

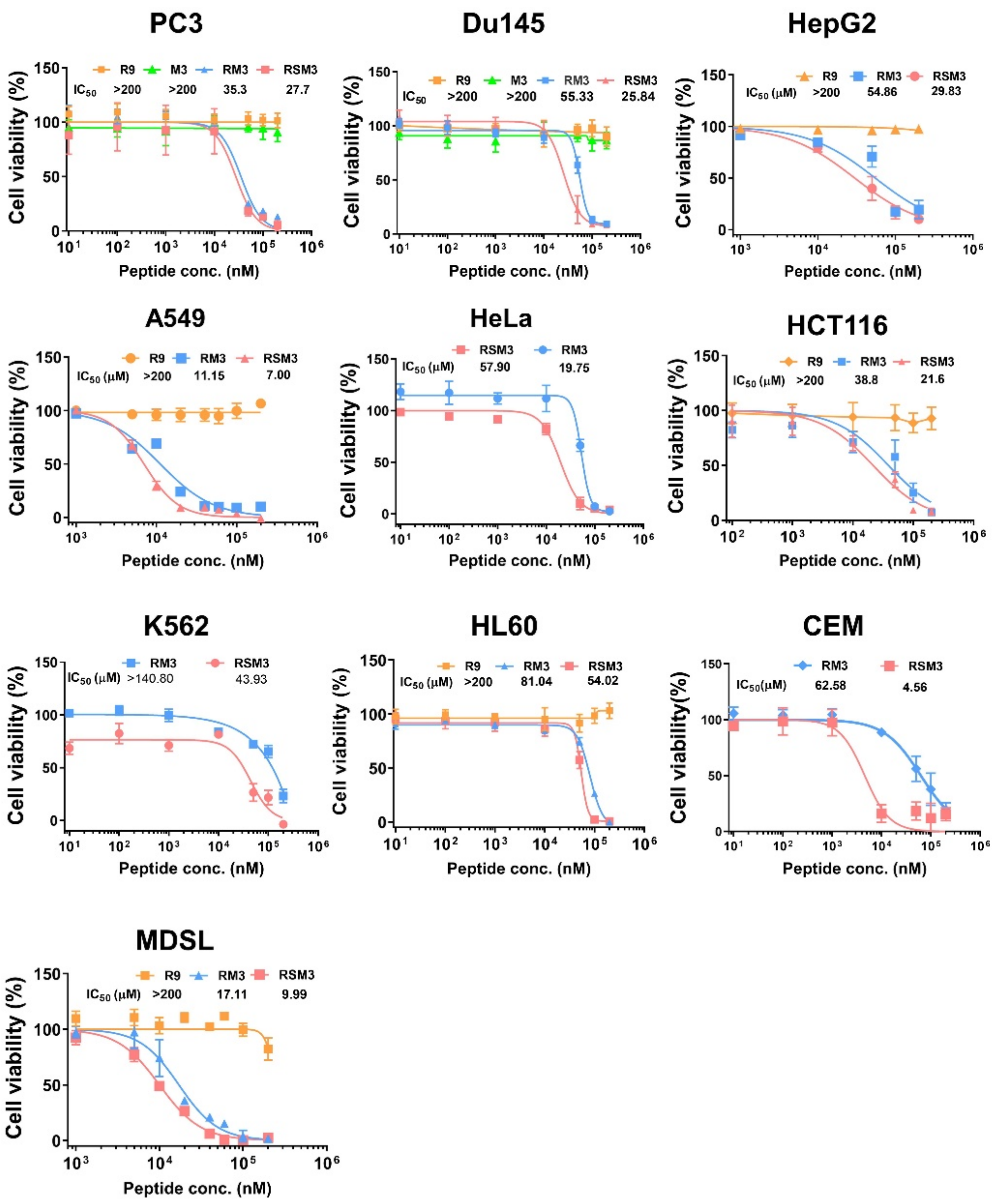


**Figure S8.** The MTT assay determinate the cytotoxicity of peptide inhibitors towards multiple cancer cell lines.

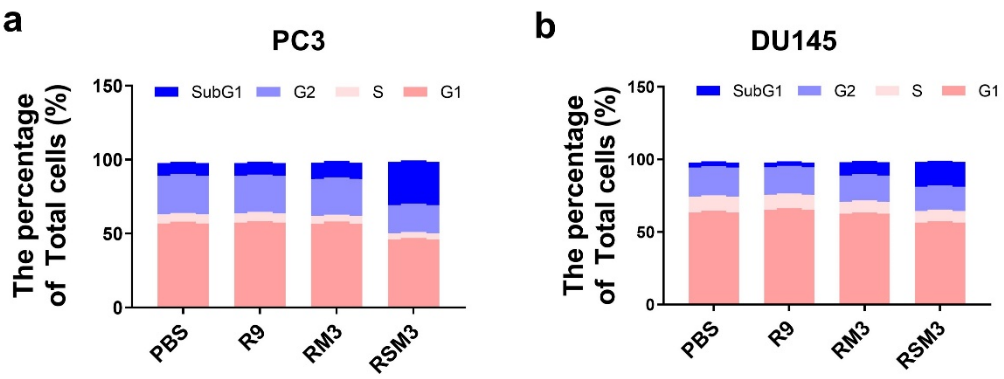


**Figure S9.** Flow cytometry investigating the effect of **RSM3** (15 μM) on the cell cycle of **(a)** PC3 and **(b)** DU145 cells over 12 h.

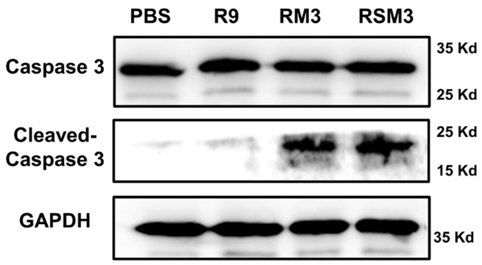


**Figure S10.** Western Blotting analysis of the activation of capase3 in PC3 cells after 15 μM peptide inhibitor treatment for 12 h.

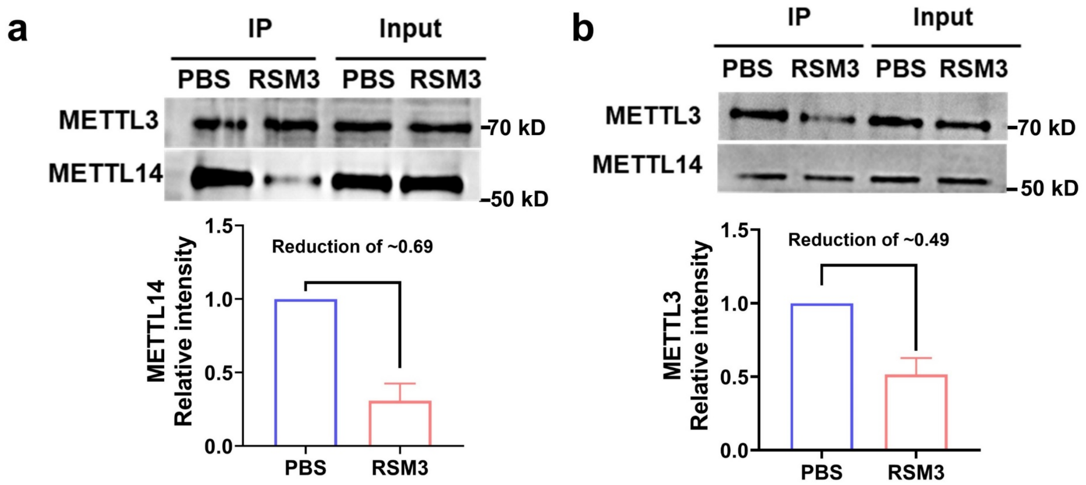


**Figure S11.** Co-IP assessing the interaction between METTL3 and METTL14 in the absence or presence of **RSM3**. The mean and error (±s.d.) were obtained from three replicates.

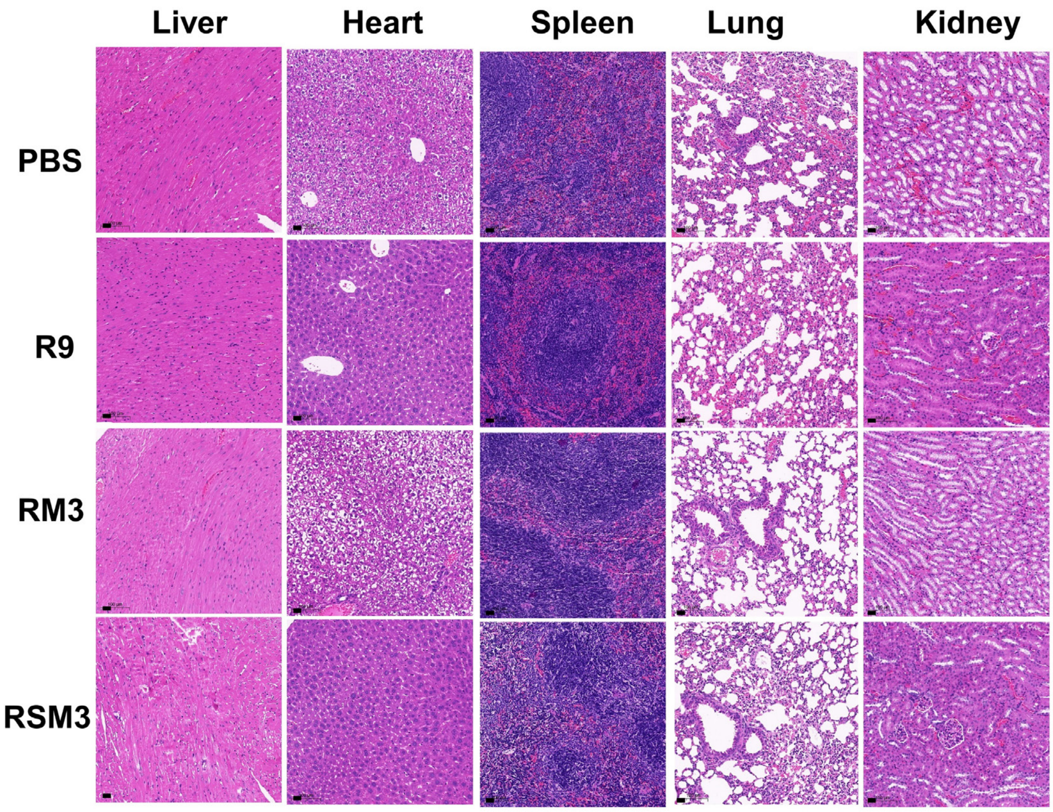


**Figure S12**. H&E staining of organ sections, including liver, heart, spleen, lung, and kidney, obtained from C57/BL mice after once 50 mg/kg peptide treatment for 24 h, scale bar 20 μm.

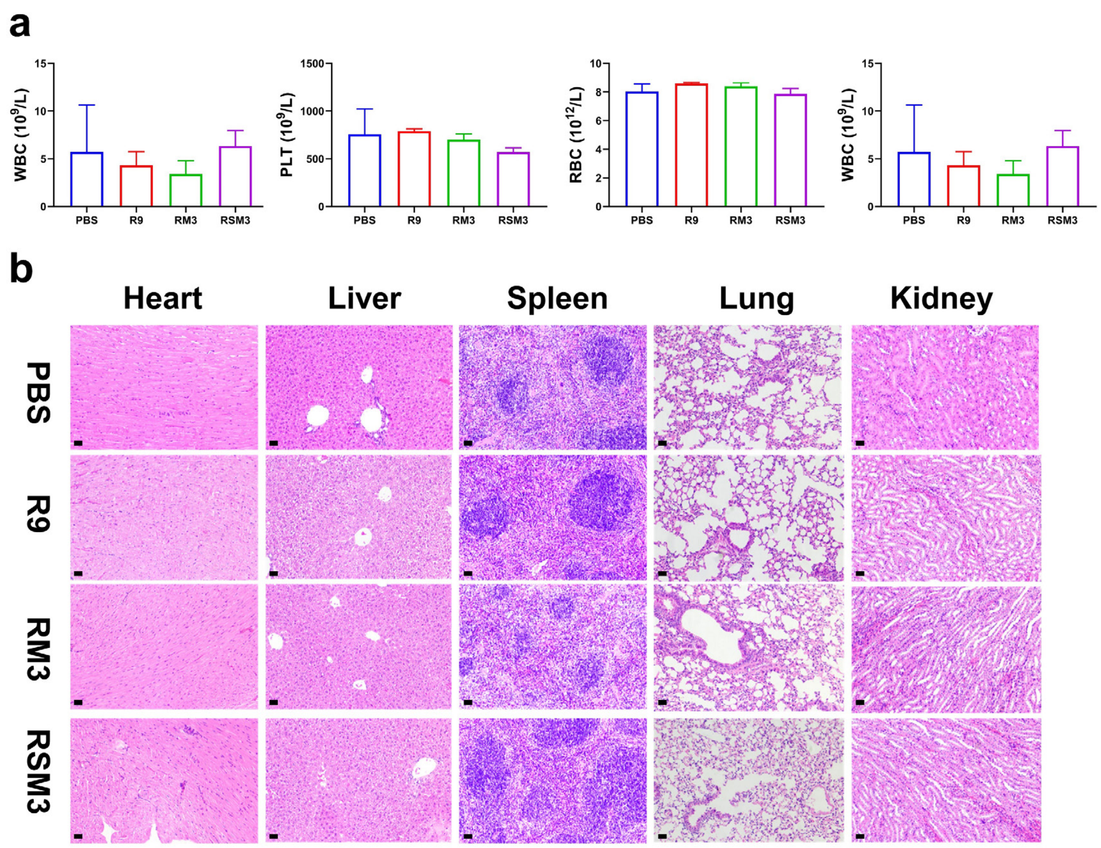


**Figure S13**. Blood routine **(a)** and H&E staining **(b)** of organ sections, including liver, heart, spleen, lung, and kidney, obtained from C57/BL mice after multiple dose injection with 50 mg/kg peptide, scale bar 20 μm.

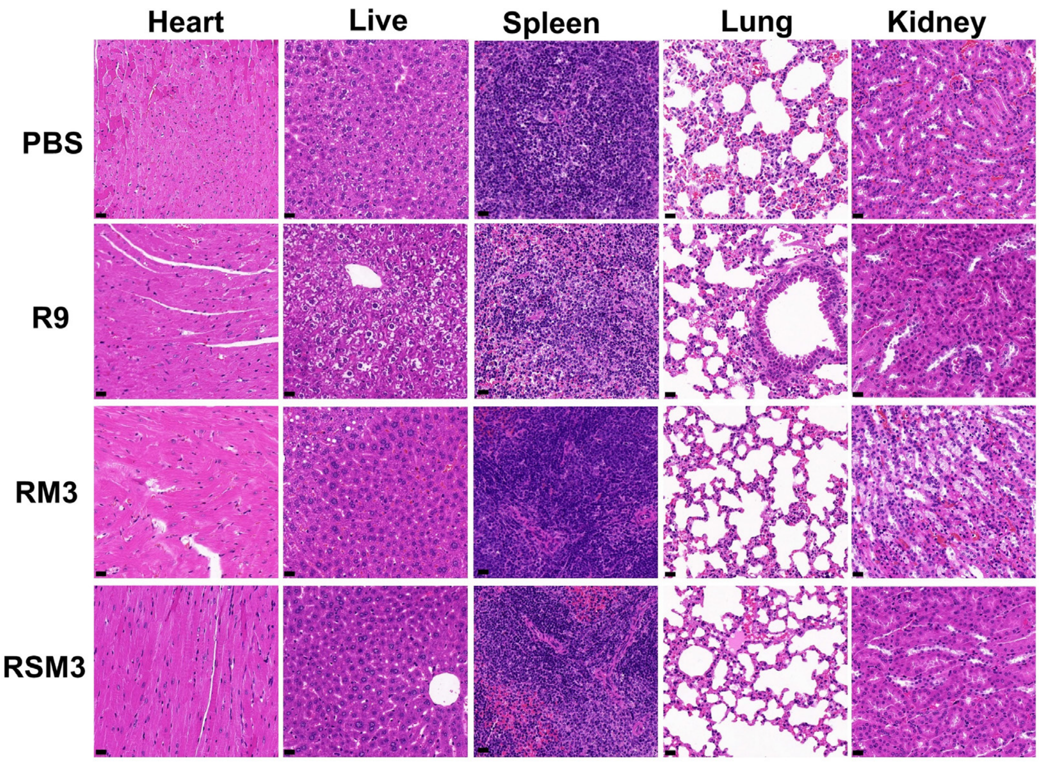


**Figure S14.** H&E staining of organ sections, including liver, heart, spleen, lung, and kidney, obtained from mice with PC3 cell xenograft tumors after 20 mg/kg peptide treatment for 10 days, scale bar 20 μm.

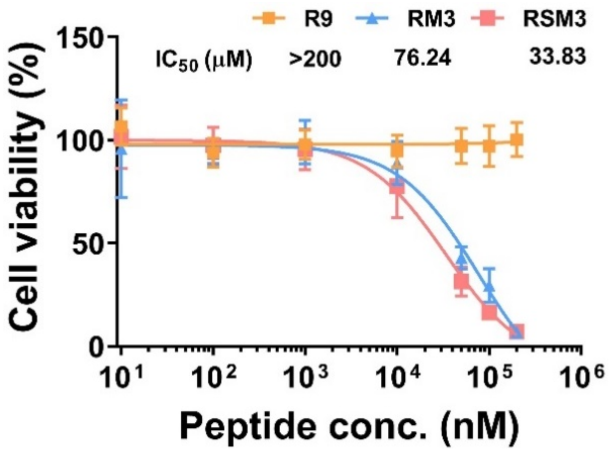


**Figure S15.** The MTT assay determinate the cytotoxicity of peptide inhibitors towards pancreatic cancer cell ASPC-1.

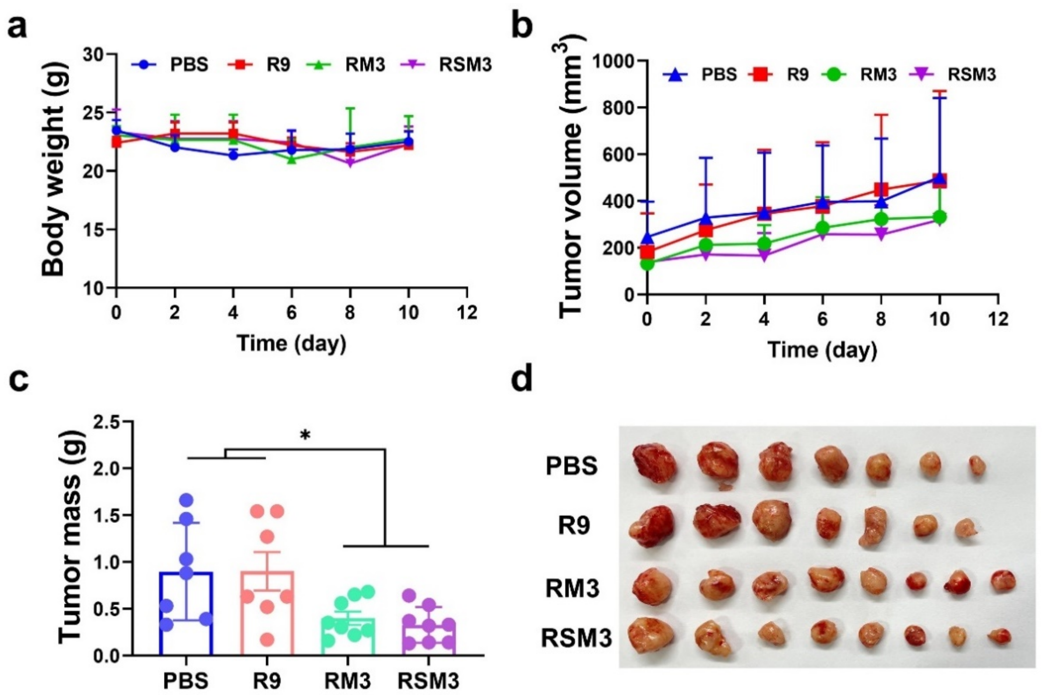


**Figure S16.** Peptide inhibitors suppress tumor growth in pancreatic cancer cell ASPC-1 xenograft tumor model. **(a)** Body weight of mice remained unchanged after treatment with peptides. **(b)** The tumor volume showed a reduction after different treatments. **(c)** Tumor mass after 10 days of treatment exhibited a notable decrease in **RM3** and **RSM3** group. **(d)** Images of sectioned tumors after 10 days of treatment displayed reduced tumor size in **RM3** and **RSM3** group. All peptides were administered at a dosage of 20 mg/kg in this assay. For **a-c** mean and error (±s.d.) were obtained from at least three replicates. One-way analysis of variance (ANOVA) with Tukey's test correction was used in statistical analysis test. $^{*}P < 0.05$, $^{**}P < 0.01$, $^{***}P < 0.005$, and not significant by $P > 0.05$.

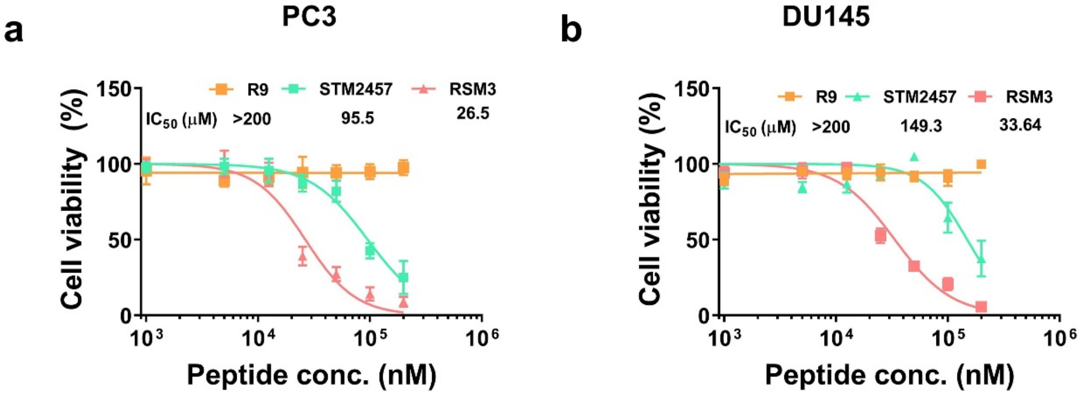


**Figure S17.** Cytotoxicity of **RSM3** compared to small molecular inhibitor **STM2457** in prostate cancer cell lines **(a)** PC3 and **(b)** DU145.

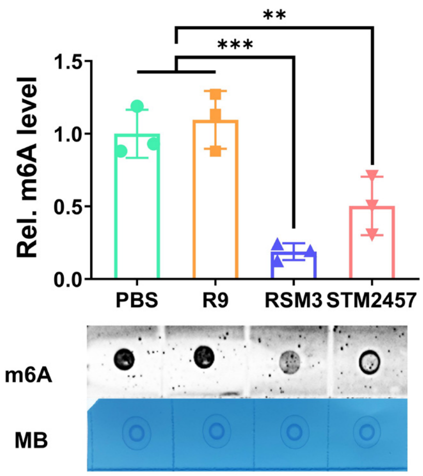


**Figure S18.** RNA blot assay(blow) and quantification (top) showing the different effects of **RSM3** and **STM2457** on m6A methylation level in PC3 cells. For mean and error (±s.d.) were obtained from three replicates. One-way analysis of variance (ANOVA) with Tukey's test correction was used in statistical analysis test. *P < 0.05, **P < 0.01, ***P < 0.005, and not significant by P > 0.05.

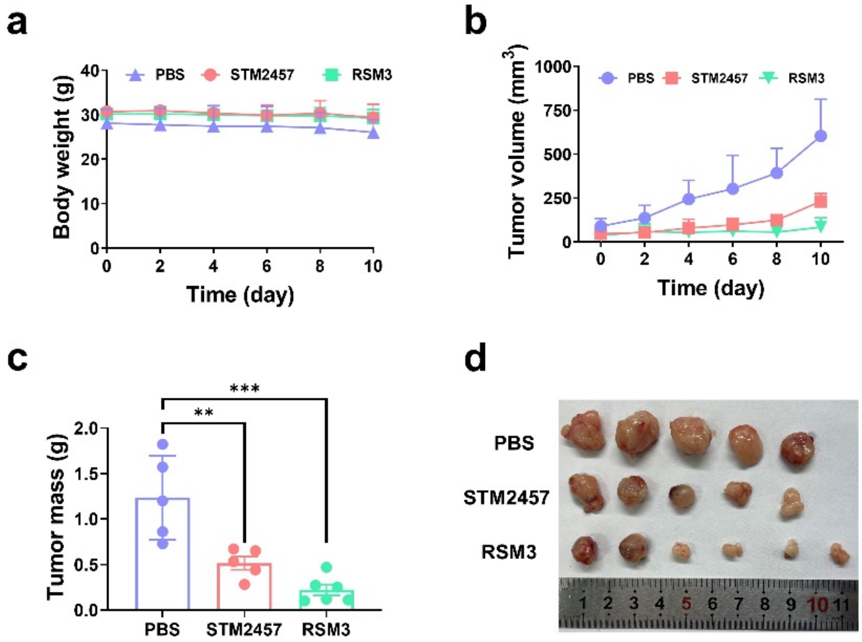


**Figure S19.** Therapeutic effect of **RSM3** compared to small molecular inhibitor **STM2457** in a prostate cancer model. **(a)** The Body weight of mice remained unchanged after treatment with both **STM2457** and **RSM3**. **(b)** The tumor volume showed a significant reduction after treatment with both inhibitors. **(c)** Tumor mass after 10 days of treatment exhibited a notable decrease in both **STM2457** and **RSM3** group. **(d)** Images of sectioned tumors after 10 days of treatment revealed reduced tumor size in **STM2457** and **RSM3** group. All molecules were administered at a dosage of 20 mg/kg in this assay. For **a-c** mean and error (±s.d.) were obtained from at least three replicates. One-way analysis of variance (ANOVA) with Tukey's test correction was used in statistical analysis test. $^*P < 0.05$, $^{**}P < 0.01$, $^{***}P < 0.005$, and not significant by $P > 0.05$.
